# Taste papilla cell differentiation requires tongue mesenchyme via ALK3-BMP signaling to regulate the production of secretory proteins

**DOI:** 10.1101/2023.04.03.535414

**Authors:** Mohamed Ishan, Zhonghou Wang, Peng Zhao, Yao Yao, Steven Stice, Lance Wells, Yuji Mishina, Hong-Xiang Liu

## Abstract

Taste papillae are specialized organs each of which is comprised of an epithelial wall hosting taste buds and a core of mesenchymal tissue. In the present study, we report that during the early stages of embryonic development, bone morphogenetic protein (BMP) signaling mediated by type 1 receptor ALK3 in the tongue mesenchyme is required for the epithelial Wnt/β-catenin activity and taste papilla cell differentiation. Mesenchyme-specific knockout (*cKO*) of *Alk3* using *Wnt1-Cre* and *Sox10-Cre* resulted in an absence of taste papillae at E12.0. Biochemical and cell differentiation analyses demonstrated that mesenchymal ALK3-BMP signaling governs the production of previously unappreciated secretory proteins, i.e., suppresses those that inhibiting and facilitates those promoting taste cell differentiation. Bulk RNA-Sequencing analysis revealed many more differentially expressed genes (DEGs) in the tongue epithelium than in the mesenchyme in *Alk3 cKO* vs control. Moreover, we detected a down-regulated epithelial Wnt/β-catenin signaling, and taste papilla development in the *Alk3 cKO* was rescued by GSK3β inhibitor LiCl, but not Wnt3a. Our findings demonstrate for the first time the requirement of tongue mesenchyme in taste papilla cell differentiation.

**Summary statement:** This is the first set of data to implicate the requirement of tongue mesenchyme in taste papilla cell differentiation.

## Introduction

Taste buds in the mammalian tongue are clusters of specialized cells that reside in taste papillae. Although structurally recognizable taste buds form later [∼E18.5 in mice (Barlow, 2015)] than taste papillae (E14.5) (Barlow, 2015, Krimm et al., 2015), taste bud cell progenitors are specified early. In mice, the tongue emerges as lingual swellings on the branchial arches at embryonic day (E) 11.0-11.5. The homogeneous epithelial cells in the primordial tongue express pan-taste cell marker Keratin 8 (Krt8) (Kramer et al., 2019, Mbiene and Roberts, 2003) and taste papilla marker sonic hedgehog (Shh) (Kramer et al., 2019, Castillo-Azofeifa et al., 2017, Liu et al., 2004, Mistretta et al., 2003). While the swellings fuse to form a spatulate tongue at E12, taste papillae appear as epithelial thickenings (papilla placodes) on the dorsal surface, and the Krt8^+^Shh^+^ cells can rapidly differentiate into two groups: taste papilla (Krt8^+^Shh^+^) and inter-papilla (Krt8^-^Shh^-^) cells. The former gives rise to taste bud cells (Thirumangalathu et al., 2009, Gaillard et al., 2015, Kramer et al., 2019) , and the latter gives rise to basal epithelial cells that remain as a progenitor population for the cell renewal of mature taste buds throughout their lifetime (Thirumangalathu et al., 2009, Miura et al., 2014, Okubo et al., 2009, Kramer et al., 2019).

There has been a well-documented general concept that cell differentiation of epithelial appendages requires mesenchymal-epithelial interactions via molecular signaling (Santosh and Jones, 2014). Unlike many other epithelial appendages, of which the cell differentiation’s regulation by the surrounding mesenchyme is well-characterized (Sennett and Rendl, 2012, Chuong et al., 2000), research on taste cell differentiation has mainly focused on signaling molecules in the epithelium (Gaillard et al., 2017, Iwatsuki et al., 2007, Liu et al., 2013, Liu et al., 2004, Mistretta et al., 2003). Less is known about the roles of the underlying tongue mesenchyme (Beites et al., 2009, Petersen et al., 2011, Prochazkova et al., 2017). Among the multiple molecular signaling pathways regulating taste papilla formation and epithelial cell differentiation, the effect of bone morphogenetic protein (BMP) pathway on the development of taste papillae are profound (Beites et al., 2009, Ishan et al., 2020, Zhou et al., 2006). However, it is unclear regarding the specifics including the mediating BMP receptor(s) of the signaling, involved tissue compartments/cell types, and interacting signaling in the epithelium.

In this study, we report that neural crest derived mesenchyme-specific conditional knockout of type I BMP receptor *Alk3* (*Alk3 cKO*), but not *Alk2 cKO*, resulted in a complete loss of taste papilla placodes (i.e., taste cell progenitors). In combination with RNA sequencing, Liquid chromatography-Mass spectrometry (LC-MS), and cell differentiation analyses using tongue organ cultures, we found that *Alk3 cKO* mesenchyme produce secreting proteins ranging 10-100-kDa that suppress the taste papilla cell differentiation through suppression of Wnt/β-catenin signaling activity. Together, our data demonstrate that BMP signaling mediated by ALK3 (ALK3-BMP signaling) in neural crest-derived mesenchyme is essential for the differentiation of taste papilla cells through suppressing the production of previously unappreciated mesenchymal secretory proteins and promoting the epithelial Wnt/β-catenin signaling activity.

## Results

### Bone morphogenetic protein (BMP) signaling is active during early taste papilla cell differentiation with the highest *Alk3* expression among type I receptors

Phosphorylation (p) of Smad1/5/8 is a crucial step and reliable indicator of BMP signaling activity (Bragdon et al., 2011, Wang et al., 2014, Massagué et al., 2005, Retting et al., 2009) . Transcripts of *Smad1*, *Smad4*, *Smad5*, and *Smad8* were detected in the tongue mesenchyme at E12.5, a stage when the tongue epithelial cells differentiate into two groups (Kramer et al., 2019): Shh^+^Krt8^+^ taste cell progenitors in taste papilla placodes and Shh^-^Krt8^-^ non-gustatory cells between papillae (Fig. 1A). At this stage, p-Smad1/5/8^+^ cells were abundantly distributed in the tongue mesenchyme and epithelium (Fig. 1B).

**Figure 1.**
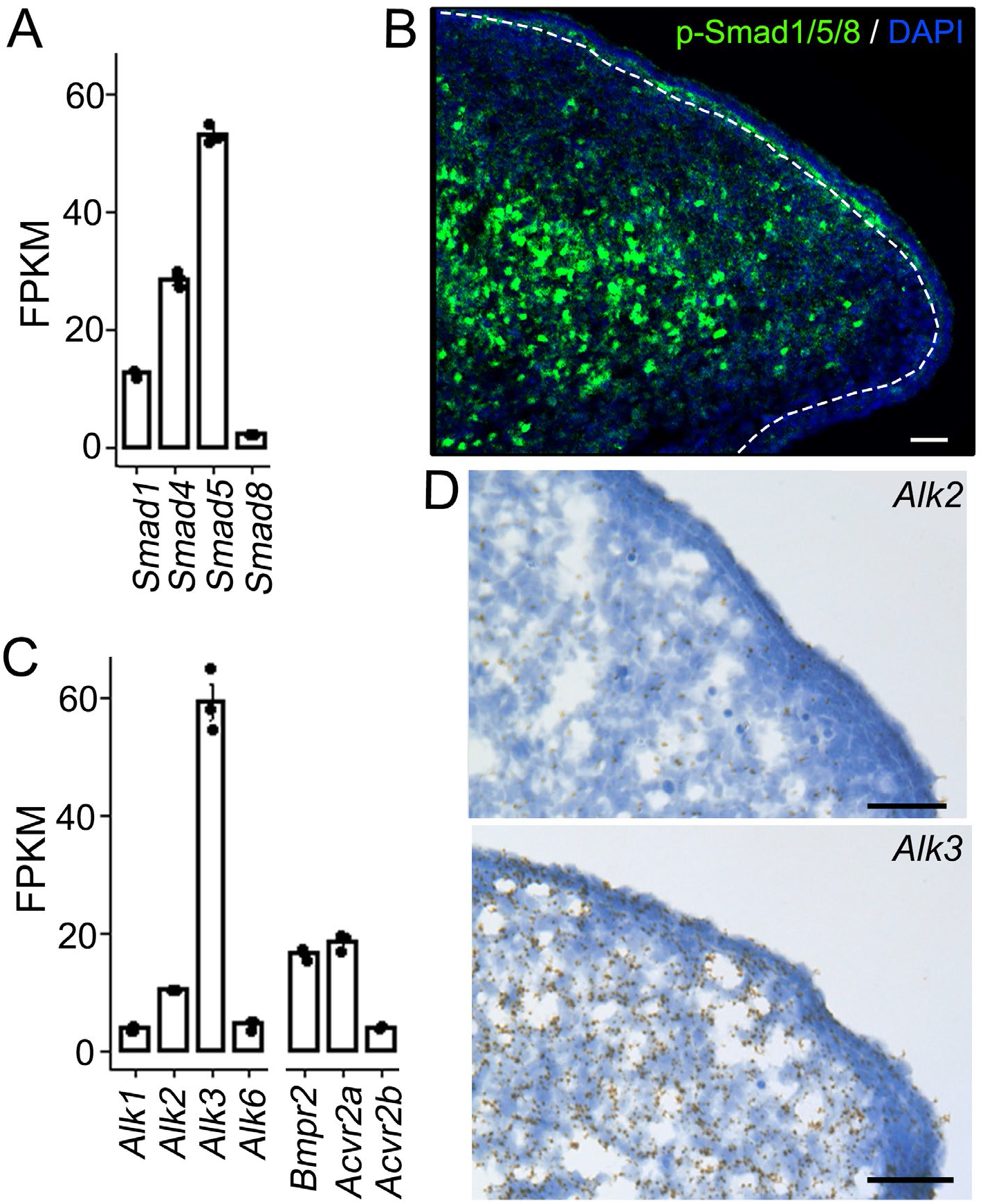
BMP signaling pathway is intact and active in E12.0-E12.5 mouse tongue tissues. **<u>A</u>**: Histograms (X±SD; n=3) to present the Fragments Per Kilobase Million (FPKM) values of BMP signaling molecules Smad1, 4, 5, 8 in the tongue mesenchyme at E12.5. **<u>B</u>**: Representative images (single-plane laser scanning confocal) of sagittal tongue sections at E12.5. The sections were immunostained for BMP signaling transcription factor p-Smad1/5/8 (green) and counterstained with DAPI (blue). Scale bar: 50 μm. **<u>C</u>**: Histograms (X±SD; n=3) to present the expression level of BMP receptors as FPKM values in the tongue mesenchyme at E12.5. **<u>D</u>**: Representative light microscopy images of E12.5 tongue sections. RNAscope *in situ* hybridization was performed using an antisense probe for *Alk2* or *Alk3* (brown). Sections were counterstained with 50% Hematoxylin to show cell nuclei (blue). Scale bars: 50 µm.

To understand the BMP signaling pathway in taste organogenesis, transcriptomic analyses revealed that among the four type I BMP receptors, the *Alk3* transcripts level was significantly higher than the other three (*Alk1*, *Alk2*, and *Alk6*) in the tongue mesenchyme (Fig. 1C) at the order of *Alk3* >> *Alk2* > *Alk6* ≈ *Alk1*. RNAscope *in situ* hybridization data further confirmed a significantly higher level of *Alk3* RNA expression than that of *Alk2* (Fig. 1D).

### Mesenchyme-specific knockout of *Alk3*, but not *Alk2*, leads to an absence of taste papillae in early embryos

To understand the role of ALK3-mediated BMP (hereafter ALK3-BMP) signaling in the tongue mesenchyme, mesenchyme-specific knockout of *Alk3* (*Alk3 cKO*) were generated using *Wnt1-Cre* (Jiang et al., 2000) and *Sox10-Cre* (Matsuoka et al., 2005) that largely labels the mesenchymal cells in the developing tongue (Liu et al., 2012, Thirumangalathu et al., 2009, Ishan et al., 2021, Yu et al., 2020). In *Wnt1-Cre/Alk3 cKO* tongues, *Alk3* transcripts were significantly reduced in the mesenchyme (Fig. 2A-B) especially in the mesenchyme immediately under the epithelium (arrowheads in Fig. 2A) compared to control. In accordance with the reduction of *Alk3* transcripts, p-Smad1/5/8^+^ cells were seen in a significantly low number in the tongue mesenchyme of E12.0 *Wnt1-Cre/Alk3 cKO* tongue (Fig. 2C, *p*<0.05 in Fig. 2D) compared to *Cre^-^/Alk3^fx/fx^* littermates (Fig. 2C). The difference was especially obvious in the mesenchymal layer immediately under the epithelium.

**Figure 2.**
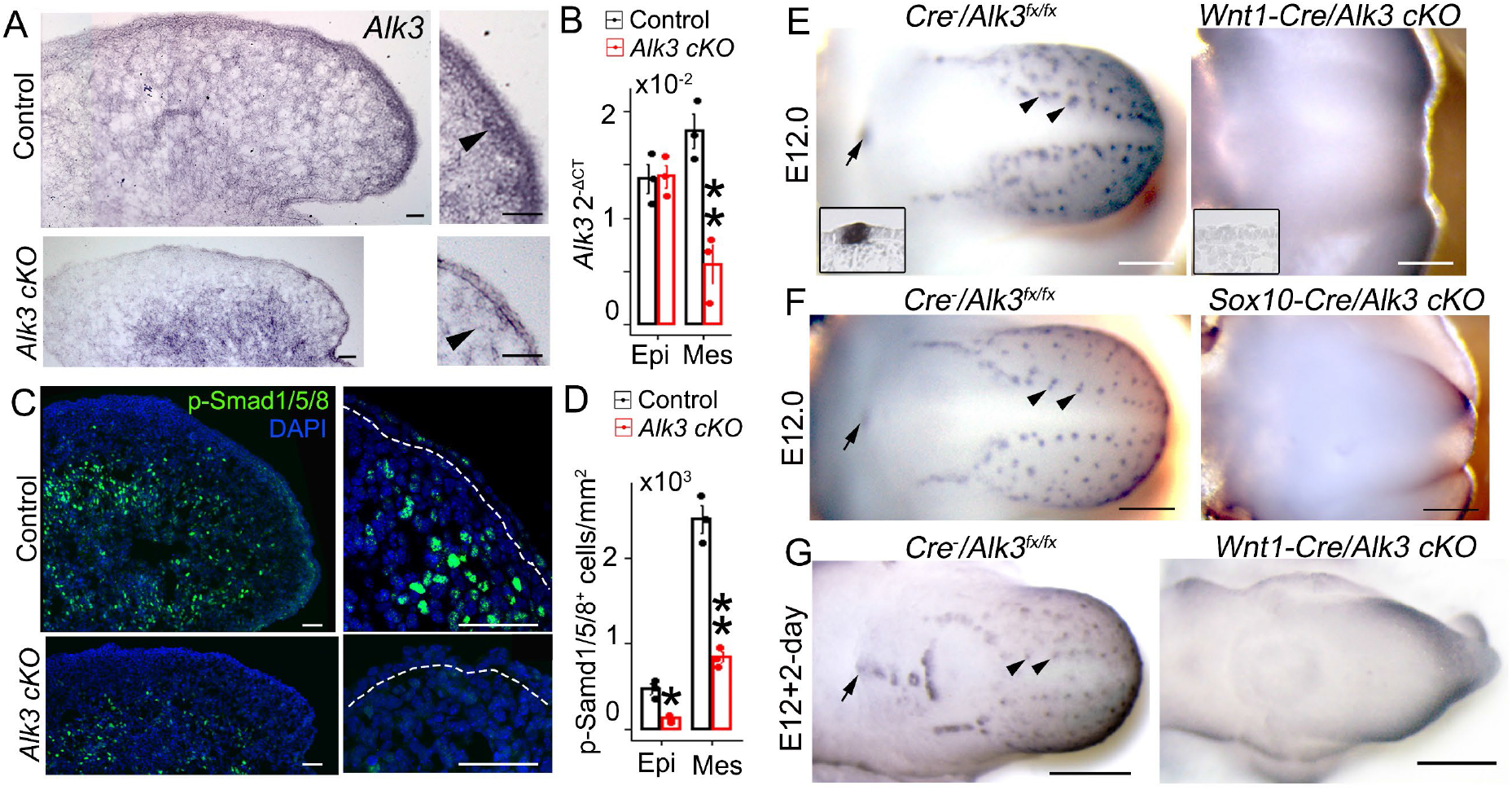
Mesenchyme-specific (*Wnt1-Cre* or *Sox10-Cre*) *Alk3 cKO* results in a complete non-gustatory epithelial cell fate. **<u>A</u>**: Light microscopy images of sagittal tongue sections of E12.0 *Cre^-^/Alk3^fx/fx^* (Control) and *Wnt1-Cre/Alk3 cKO* (*Alk3 cKO*) mice. *In situ* hybridization was performed using an antisense probe for *Alk3* (purple). Arrowheads point to the mesenchymal region immediately under the epithelium. Scale bars: 100 µm. **<u>B</u>**: A histogram (X±SD; n=3) to present the 2^-ΔCT^ values of *Alk3* gene transcripts in the tongue epithelium (Epi) and mesenchyme (Mes) of *Cre^-^/Alk3^fx/fx^* and *Wnt1-Cre/Alk3 cKO* mouse embryos at E12.0. \*\**p*≤0.01 compared to *Cre^-^/Alk3^fx/fx^* littermates using two-way ANOVA followed by Fisher’s least significant difference (LSD) analyses. **<u>C</u>**: Single-plane laser scanning confocal images of sagittal tongue sections that were immunostained for p-Smad1/5/8 (green) in *Cre^-^/Alk3^fx/fx^* (Control) and *Wnt1- Cre/Alk3 cKO* (*Alk3 cKO*) mice. The right column illustrates the anterior tongue region at a high magnification. Scale bars: 50 μm. **<u>D</u>**: A histogram (X±SD; n=3) to present the number of p-Smad1/5/8^+^ cells per mm^2^ in the tongue epithelium (Epi) and mesenchyme (Mes) of *Cre^-^/Alk3^fx/fx^* (Control) and *Wnt1-Cre/Alk3 cKO* (*Alk3 cKO*) mice. \**p*≤0.05, \*\**p*≤0.01 compared to *Cre^-^/Alk3^fx/fx^* littermate controls using two-way ANOVA followed by Fisher’s LSD analyses. **<u>E-G</u>**: Representative light microscopy images of E12.0 tongues immunostained for the developing taste papilla marker sonic hedgehog (Shh) (blue). Insets in E are light microscopy images of sagittal tongue sections immunostained for Shh. Arrowheads and arrows point to Shh^+^ fungiform and circumvallate papilla placodes respectively. Scale bars: 200 µm.

A dramatic phenotype of mesenchyme-specific deletion of ALK3-BMP signaling is a complete loss of taste papillae in E12.0 *Wnt1-Cre/Alk3 cKO* (Fig. 2E). Such a complete loss of taste papilla cells was consistently observed (Fig. 2F) in *Alk3 cKO* using another *Cre* driver, *Sox10-Cre* that exclusively marks neural crest-derived mesenchyme (Yu et al., 2020) (Supplemental Fig. 1). In addition, a truncation of the tip of tongue/mandible was observed in both *Wnt1-Cre/Alk3 cKO* and *Sox10-Cre/Alk3 cKO* mice (Fig. 2E-F). Of note, mesenchyme-specific knockout of another BMP receptor *Alk2* did not lead to an absence of taste papillae (Supplemental Fig. 2); instead, well-developed and stereotypically located fungiform (arrowheads in Supplemental Fig. 2) and circumvallate (the arrow in Supplemental Fig. 2) papilla placodes were seen in the *Wnt1-Cre/Alk2 cKO* similarly to the *Cre^-^* littermates (Fig. 2E-F, Supplemental Fig. 2).

Due to the embryonic lethality of *Alk3* c*KO* driven by *Wnt1-Cre* (sudden death due to heart failure at E12.5 or later) (Stottmann et al., 2004), close attention was paid to the condition of all embryos at collection (E12 or younger) to ensure the rigorous heart beating and blood circulation. Cell viability was evaluated using cell proliferation and apoptosis markers including Ki67^+^ (pan proliferation), BrdU^+^ (S-phase), p-H3^+^ (M-phase), and cleaved (c)-Caspase3^+^ (c-Cas3, apoptosis). No changes in cell proliferation and apoptosis (Supplemental Fig. 3) were detected in mesenchymal *Alk3 cKO* (*Wnt1-Cre* and *Sox10-Cre*) tongues compared to that of *Cre^-^/Alk3^fx/fx^* littermates.

To understand the developmental course of taste papilla absence in E12.0 mesenchymal *Alk3 cKO*, phenotypes were analyzed in earlier stages (E10.5-E11.5) of embryos and further developed tongue organs in the 2-day cultures started at E12.0. Alterations of tongue development was not obvious in *Alk3 cKO* until E11.5 at which the lateral tongue swellings were smaller in *Alk3 cKO* than littermate controls with the same somite numbers (Supplemental Fig. 4). The four branchial arches (E10.5) and early tongue swellings (E11.0) developed in *Wnt1- Cre/Alk3 cKO* mice similarly to *Cre^-^/Alk3^fx/fx^* littermates (Supplemental Fig. 4). After 2 days extension of E12 tongues in culture, taste papillae did not develop in *Wnt1-Cre/Alk3 cKO* (Fig. 2G). Noteworthy, the cultured *Alk3 cKO* tongues that were free from the restraint of mandible depicted apparent outgrowth and a pointed tip (Fig. 2G).

To understand whether the elevated level of ALK3-BMP signaling alters taste papilla cell differentiation, *Wnt1-Cre* mediated constitutively activated (ca) *Alk3* (*Wnt1-Cre/caAlk3*) was analyzed. No obvious changes in the taste papilla and taste bud development were found (Supplemental Fig. 5). Similar to the *Cre^-^* littermate controls, fungiform and circumvallate papillae (E12.5, E14.5) and taste buds (postnatal day 21) developed in the *Wnt1-Cre/caAlk3* mice (Supplemental Fig. 5).

### *Alk3 cKO* mesenchyme-derived proteins result in an absence of taste papillae mimicking the *in vivo* phenotype of *Alk3 cKO*

Tissue-tissue and cell-cell interactions may be through direct contact (Cutler and Gremski, 1991) and/or secretory signals (Nakano et al., 2017, Thesleff et al., 1988, Sagmeister et al., 2008, Horowitz and Thannickal, 2006, Chapman, 2011, Rice et al., 2004, Prochazkova et al., 2017). To understand how mesenchymal ALK3-BMP interact with overlying epithelium to regulate taste papilla cell differentiation, tongue mesenchyme tissue of E12.0 *Wnt1-Cre/Alk3 cKO* or *Cre^-^/Alk3^fx/fx^* was placed adjacently to E12.0 wild type (WT) tongues for 2-day cultures (Fig. 3A, B). In the WT tongues cultured with *Wnt1-Cre/Alk3 cKO* tongue mesenchyme, Shh^+^ taste papillae were significantly reduced in number and less profound compared to those co-cultured with control tongue mesenchyme (Fig. 3A, *p*<0.01). Furthermore, we collected the medium from tongue mesenchymal cell cultures (mesenchyme-conditioned medium) and tested the effects. Mesenchyme-conditioned medium (Mes-CM) from *Wnt1-Cre/Alk3 cKO* tongues showed potent inhibitory effects on taste papilla cell differentiation in WT tongues compared to control group (Fig. 3B, *p*<0.01). The morphology and proliferating rate of the tongue mesenchymal cells were similar in *Alk3 cKO* and control groups (Supplemental Fig. 6).

**Figure 3.**
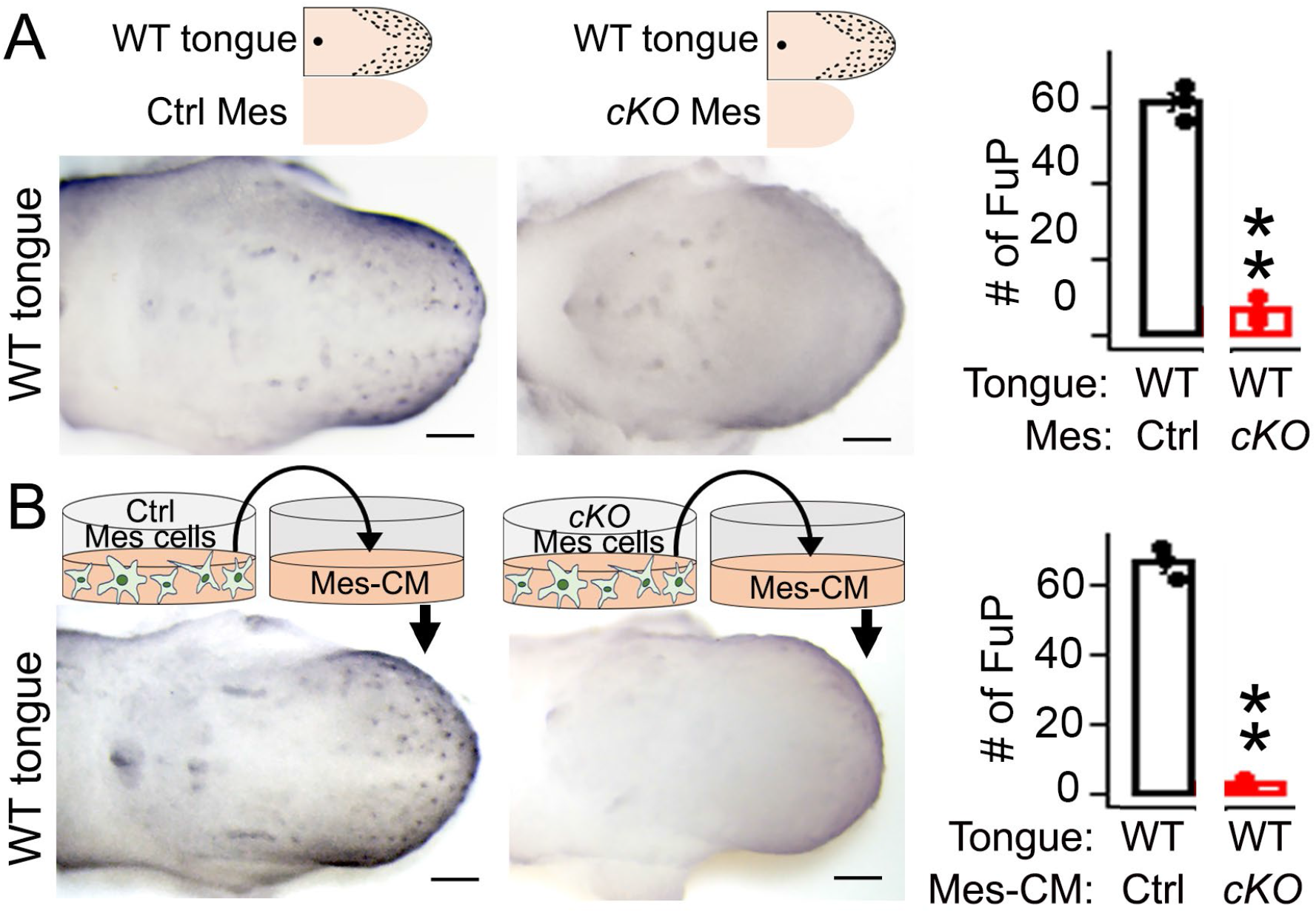
Differentiation of taste papilla cells is suppressed by *Wnt1-Cre/Alk3 cKO* tongue mesenchyme and conditioned medium in E12.0+2-day wild type tongue cultures. **<u>A-B</u>:** Representative light microscopy images of E12.0+2-day tongue cultures from wild type (WT) mice. Cultures were either co-cultured with tongue mesenchyme (Mes) (A) or fed with mesenchyme-conditioned medium (Mes-CM) (B) and immunostained for Shh (blue). Ctrl: *Cre^-^/Alk3^fx/fx^*; *cKO*: *Wnt1-Cre/Alk3 cKO*. Scale bars: 200 µm. Histograms (X±SD; n=3) on the right show the number of Shh^+^ fungiform papillae (FuP). Schematic diagrams show how the experiment was set up. \*\**p*≤0.01 compared to *Cre^-^/Alk3^fx/fx^* littermate control using two-way ANOVA followed by Fisher’s least significant difference (LSD) analyses.

To define the tongue mesenchyme-derived factors that mediate the impact on taste papilla cell differentiation, proteins were extracted from Mes-CM, and those from *Wnt1-Cre/Alk3 cKO* tongues showed an almost complete suppression on taste papilla formation compared to control (Fig. 4B vs 4A, *p*<0.01 in 4E). Proteinase K (ProK) pretreatment efficiently digested the extracted proteins (Supplemental Fig. 7) and eliminated the inhibition of proteins from *Alk3 cKO* tongue mesenchyme (Fig. 4D vs 4C, *p*>0.05 in 4E).

**Figure 4.**
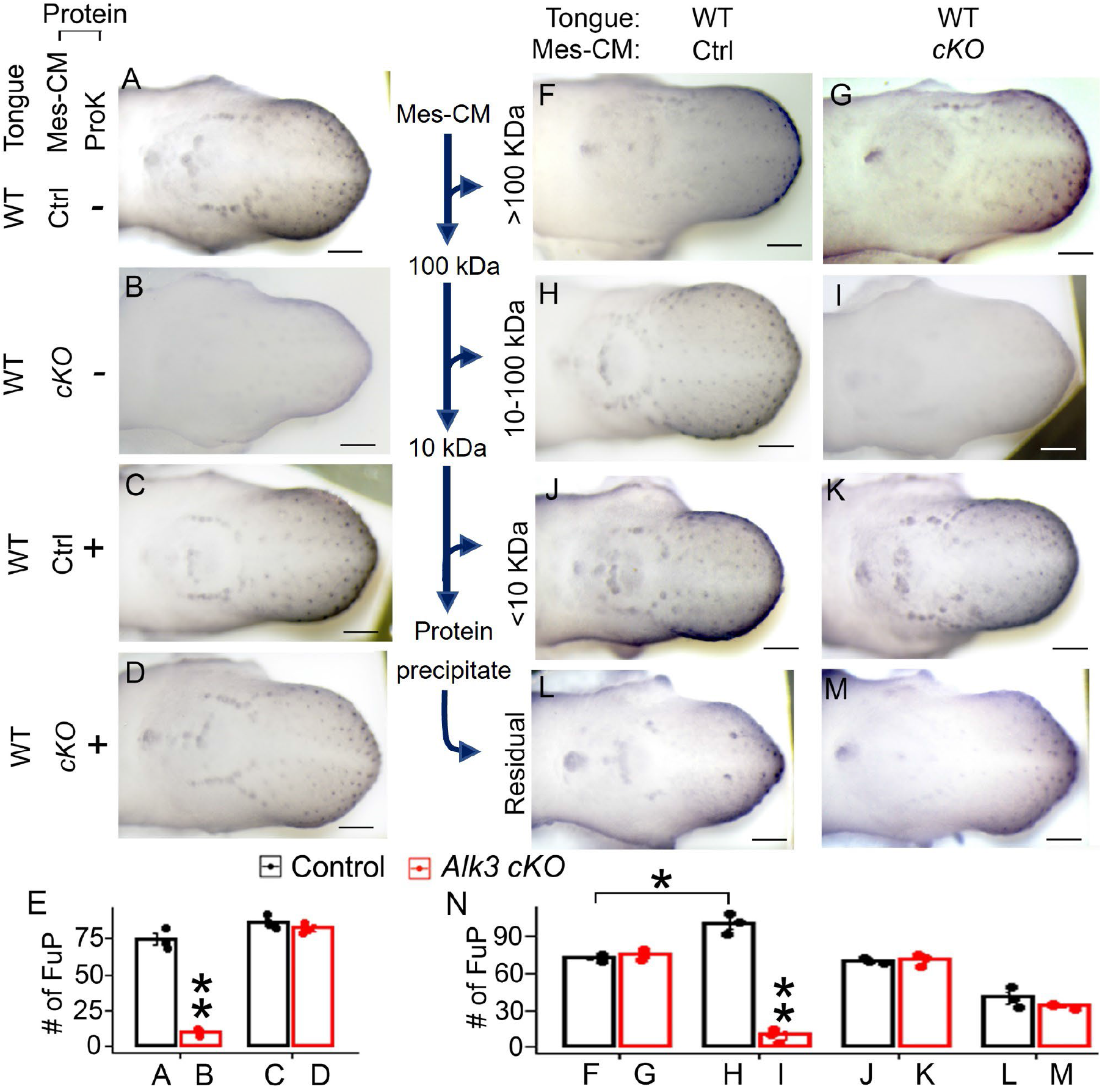
Proteins from *Alk3 cKO* tongue mesenchyme-conditioned medium inhibit the differentiation of taste papilla cells. **<u>A-D, F-M</u>**: Representative light microscopy images of E12.0+2-day WT tongues cultures immunostained for taste papilla marker Shh (blue). Proteins from mesenchyme-conditioned medium (Mes-CM) (A-D; F-K) or protein-free residual solution (L-M) were added to the culture medium. Ctrl: *Cre^-^/Alk3^fx/fx^* (A, C, F, H, J, L); *cKO*: *Wnt1- Cre/Alk3 cKO* (B, D, G, I, K, M). Scale bars: 200 µm. **<u>E, N</u>**: Histograms (X±SD; n=3) to present the number of Shh^+^ fungiform papillae (FuP) in the tongue cultures under different experimental conditions shown in A-D, F-M. \**p*≤0.05, \*\**p*≤0.01 compared to the *Cre^-^/Alk3^fx/fx^* littermate control using two-way ANOVA followed by Fisher’s LSD analyses.

The effects of mesenchyme-derived proteins were further analyzed using filtered protein fragments at different molecular weights. Administration of 10-100-kDa proteins from control group to the WT tongue cultures resulted in an increase of taste papilla number (Fig. 4H vs 4F, J, L, *p*<0.05 in Fig. 4N), while the 10-100-kDa proteins from *Alk3 cKO* lead to an absence of taste papillae (Fig. 4I vs G, K, M, *p*<0.01 in Fig. 4N) which mimics the *in vivo* phenotype (Fig. 2E). In contrast, >100-kDa or <10-kDa Mes-CM proteins or protein-free residual solution did not result in a difference of taste papillae between *Wnt1-Cre/Alk3 cKO* and control groups (Fig. 4G vs F, K vs J, M vs L, *p*>0.05 in Fig. 4N)

### Mesenchymal *Alk3 cKO* leads to a down-regulation of epithelial Wnt/β-catenin signaling

To understand the molecular basis of the potent suppression of *Alk3 cKO* on taste papilla cell differentiation, RNA-Sequencing analyses was performed on the separated epithelium and mesenchyme of E12.0 *Wnt1-Cre/Alk3 cKO* vs littermate control (*Cre^-^/Alk3^fx/fx^*) tongues. To our surprise, many more differentially expressed genes (DEGs) were detected in the epithelium than in the mesenchyme (|FC|>1, p<0.05, FDR q<0.05) (Fig. 5A). Among the total 350 DEGs, 287 genes (183 up-regulated, 104 down-regulated) were detected in the epithelium, and 58 genes (30 up-regulated, 28 down-regulated) in the mesenchyme, and 5 genes were upregulated in both the epithelium and mesenchyme (Fig. 5A, Supplemental Table 1).

**Figure 5.**
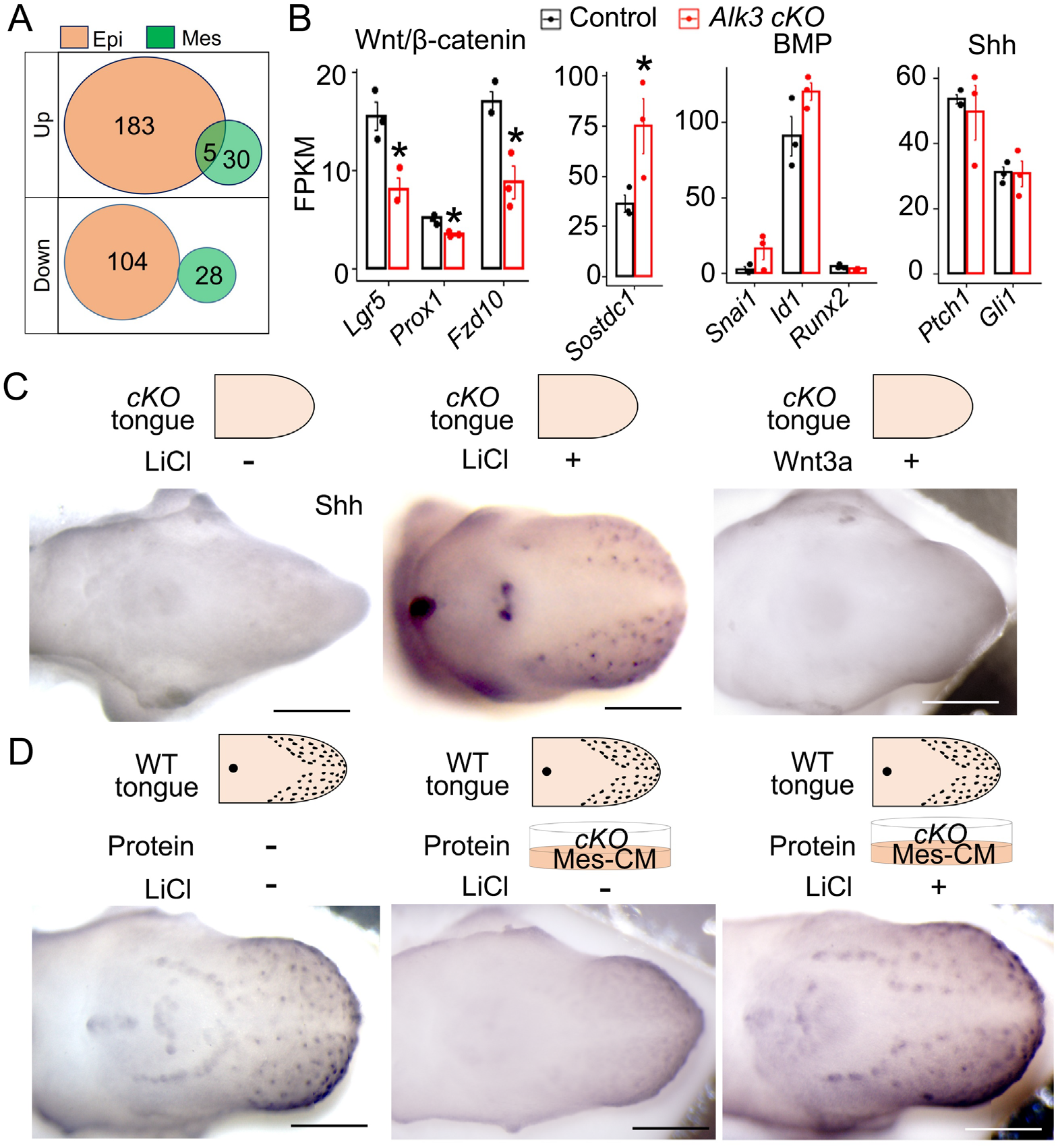
Mesenchymal *Alk3 cKO* leads to the down-regulation of Wnt/β-catenin signaling in the tongue epithelium and activating the pathway rescues taste papilla development. **<u>A</u>:** A Venn diagram to show the number (shown in the pies) of differentially expressed genes (DEGs) in the tongue epithelium (Epi) and mesenchyme (Mes) of *Wnt1-Cre/Alk3 cKO* versus *Cre^-^/Alk3^fx/fx^* littermate control. Up: Upregulated DEGs; Down: Downregulated DEGs. **<u>B</u>:** Histograms (X±SD; n=3) to present the FPKM values of signaling components in E12.0 tongue epithelium including Wnt/β-catenin, BMP, and Shh pathways. Control: *Cre^-^/Alk3^fx/fx^*; *Alk3 cKO*: *Wnt1-Cre/Alk3 cKO*. \**p*≤0.05 (adjusted p-value) DESeq2 statistical analysis based on read counts. **<u>C-D</u>:** Representative light microscopy images of E12+2-day tongue cultures from *Wnt1-Cre/Alk3 cKO* (*cKO* in C, D), or WT (D) mice. Tongue cultures were administered with 5 mM LiCl or 20% Wnt3a conditioned medium to activate Wnt/β-catenin signaling activity and immunostained for Shh (blue). Schematic diagrams show how the experiment was set up. Mes-CM: Mesenchyme-conditioned medium. Scale bars: 200 µm.

Further analyses indicated a down-regulation of Wnt/β-catenin signaling in the tongue epithelium of *Wnt1-Cre/Alk3 cKO* vs control. The epithelial Wnt/β-catenin-related DEGs include downregulated positive regulators *Lgr5*, *Prox1, Fzd10*, and the upregulated inhibitor *Sostdc1* (Fig. 5B). In contrast, known target genes of BMP or Shh singling pathways were unaltered in the epithelium of *Wnt1-Cre/Alk3 cKO* vs control (Fig. 5B).

To understand whether the downregulation of Wnt/β-catenin signaling is the main cause of absence of taste papillae caused by mesenchymal *Alk3 cKO*, Wnt/β-catenin activators LiCl or (Clément-Lacroix et al., 2005, Iwatsuki et al., 2007) Wnt3a were added to the culture medium. As predicted, control group tongues depicted an increase in taste papilla development in the E12+2-day cultures with LiCl and Wnt3a (Supplemental Fig. 8). Importantly, taste papilla development in *Wnt1-Cre/Alk3 cKO* tongue cultures was only rescued when treated with LiCl but not with Wnt3a (Fig. 5C). Moreover, the administration of LiCl also prevented *Alk3 cKO* tongue mesenchyme-derived proteins from the inhibition of taste papilla development (Fig. 5D).

### Mesenchymal *Alk3 cKO* alters the production of previously unappreciated secretory proteins

To identify the tongue mesenchyme-derived proteins that regulate epithelial cell differentiation, we compared the transcript levels of proteins-encoding genes in the tongue mesenchyme and proteomic profiles of mesenchyme-conditioned medium between *Alk3 cKO* vs control. Kyoto Encyclopedia of Genes and Genomes (KEGG) and Gene ontology (GO) analyses of RNA-Seq data revealed that the 30 up-regulated and 28 down-regulated genes in E12.0 *Alk3 cKO* mesenchyme are associated with multiple biological processes. Of note, enrichment of DEGs in protein trafficking including intracellular transport and exocytosis (Fig. 6A) was found in addition to other development-related processes. Noteworthy, transcripts level of Wnt/β-catenin signaling regulators including agonists (Wnt ligands, R-Spondins) and antagonists (Dkk’s, Sfrp’s, Sostdc1, Wif1) (MacDonald et al., 2009) were unaltered in the E12.0 *Wnt1-Cre/Alk3 cKO* mesenchyme (Fig. 6B).

**Figure 6.**
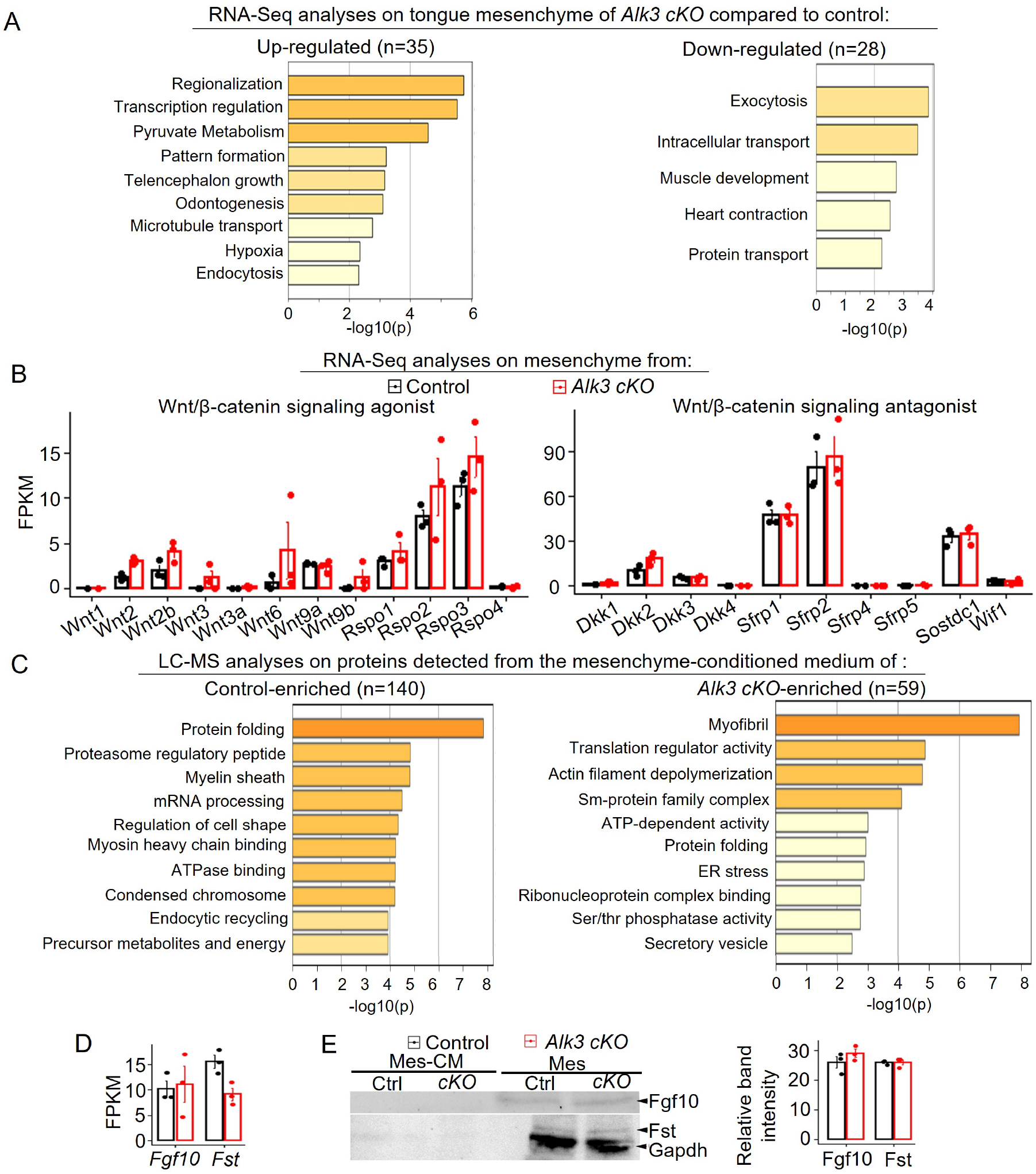
Mesenchymal *Alk3 cKO* alters the production of proteins that are not previously unappreciated. **<u>A</u>**: GO enrichment and KEGG (Kyoto Encyclopedia of Genes and Genomes) pathway analyses to present the functional associations of RNA-Seq-detected DEGs in the tongue mesenchyme of E12.0 *Wnt1-Cre/Alk3 cKO* (*Alk3 cKO*) compared to *Cre^-^/Alk3^fx/fx^* littermates. **<u>B</u>**: Histograms (X±SD; n=3) to present the FPKM values of the known regulators of Wnt/β-catenin signaling in the E12.0 tongue mesenchyme. Control: *Cre^-^/Alk3^fx/fx^*; *Alk3 cKO*: *Wnt1-Cre/Alk3 cKO*. No statistically significant differences were found with (adjusted p-value) DESeq2 statistical analysis based on read counts. **<u>C</u>**: Gene Ontology (GO) enrichment and pathway analyses to show the functional association of differentially expressed proteins in tongue mesenchyme-conditioned medium from *Wnt1-Cre/Alk3 cKO* (*Alk3 cKO*) and *Cre^-^/Alk3^fx/fx^* (Control). These extracted proteins were detected with liquid chromatography-mass spectrometry (LC-MS). **<u>D:</u>** A histogram (X±SD; n=3) to present the transcripts levels (FPKM values) of FGF10/*Fgf10*, and Follistatin/*Fst* in the tongue mesenchyme of *Cre^-^/Alk3^fx/fx^* (Control) and *Wnt1-Cre/Alk3 cKO* (*Alk3 cKO*) mice. **<u>E</u>**: Western blot bands of FGF10, Fst, and Gapdh in the mesenchyme-conditioned medium (Mes-CM) or mesenchyme (Mes) tissue from E12.0 *Cre^-^/Alk3^fx/fx^* (Ctrl) and *Wnt1-Cre/Alk3 cKO* (*cKO*). The histogram to the right (X±SD; n=3) presents the normalized band intensities of FGF10, and Fst relative to the Gapdh in E12.0 *Cre^-^/Alk3^fx/fx^* (Control) and *Wnt1-Cre/Alk3 cKO* (*Alk3 cKO*) tongue mesenchyme. No statistically significant differences were found in *Alk3 cKO* compared to the *Cre^-^/Alk3^fx/fx^* littermate control using Student’s *t*-test.

Out of the LC-MS-detected proteins in the mesenchyme-conditioned medium, 199 proteins were differentially expressed (control or *Alk3 cKO* only). The proteomic profiles of mesenchyme-conditioned media were largely altered by mesenchymal *Alk3* cKO, among which 140 were enriched in *Cre^-^/Alk3^fx/ fx^* and 59 in *Wnt1-Cre/Alk3 cKO* group (Fig. 6C). GO analysis revealed that detected proteins in *Wnt1-Cre/Alk3 cKO* group are mainly involved in post-transcriptional processes, including translation regulatory activity, post-translational modification such as regulation of endoplasmic reticulum stress, protein folding, and secretion including secretory vesicles (Fig. 6C). Noteworthy, none of the Wnt/β-catenin signaling agonist or antagonist were detected by the LC-MS. Moreover, two well-known mesenchymal secretory proteins (i.e. FGF10, and Follistatin/Fst) that are important for taste papilla development (Beites et al., 2009, Petersen et al., 2011, Prochazkova et al., 2017) appeared to be unaltered at both mRNA and proteins levels in the E12.0 *Wnt1-Cre/Alk3 cKO* mesenchyme compared to the littermate controls (Fig. 6D, E).

## Discussion

Our present study provides data demonstrating for the first time a requirement of tongue mesenchyme via ALK3-BMP signaling for the activity of Wnt/β-catenin pathway in the epithelium and taste papilla cell differentiation during early embryonic development. We propose a working model regarding how mesenchymal ALK3-BMP signaling interacts with epithelial Wnt/β-catenin pathway to regulate taste papilla cell differentiation (Fig. 7). Under normal conditions, mesenchymal ALK3-BMP signaling facilitates the secretion of proteins that promote taste papilla cell differentiation and suppresses those inhibitory proteins, thus allowing for a proper activity of epithelial Wnt/β-catenin signaling and taste papilla development. Thus, in the absence of mesenchymal ALK3-BMP signaling, the production of inhibitory secretory proteins is enhanced which causes the deficiencies of epithelial Wnt/β-catenin signaling upstream to GSK3β, thereby resulting in the absence of taste papillae.

**Figure 7.**
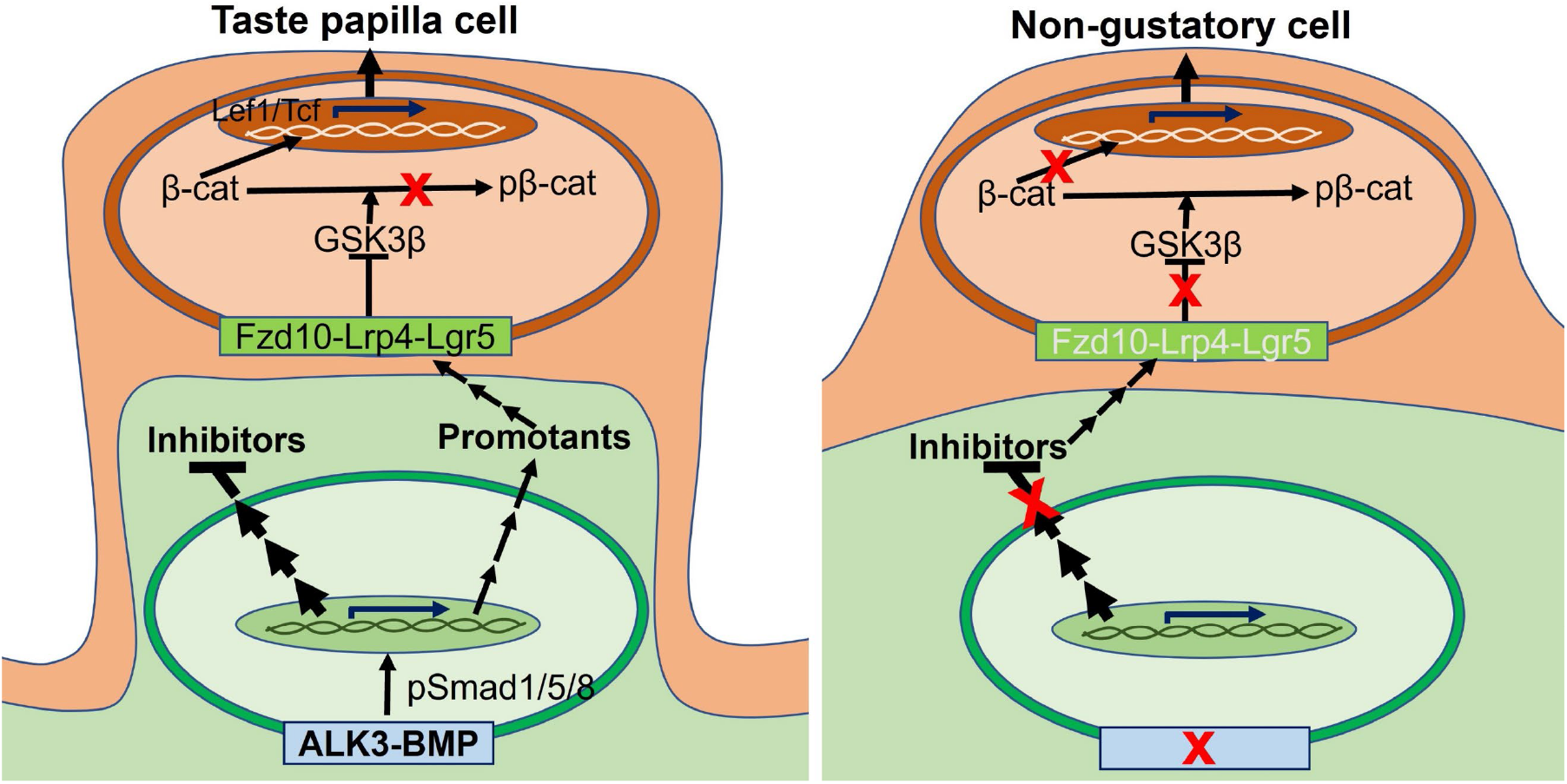
Schematic diagram illustrates a proposed model of how mesenchymal ALK3-BMP interacts with epithelial Wnt/β-catenin signaling for the differentiation of taste papilla cells. We propose that ALK3-BMP signaling in the tongue mesenchyme regulates the production of secretory proteins that promote or inhibit epithelial cell differentiation to taste papilla cells. Under normal conditions, mesenchymal ALK3-BMP signaling enhances the secretion of promotants and inhibit the secretion of inhibitors that alter the components upstream to GSK3β in the destruction complex and thus, promote or inhibit epithelial Wnt/β- catenin signaling activity.

### Tongue mesenchyme determines epithelial cell fate at early stages of taste papilla development through the production of secretory proteins that are previously unappreciated

The importance of mesenchymal-epithelial interactions in the cell differentiation of epithelial appendages has been a well-established during the development of many organs, including skin (Yamaguchi et al., 2004), mammary gland (Cunha and Hom, 1996), lung (Horowitz and Thannickal, 2006) , kidney (Müller et al., 1997), urogenital buds (Jerman et al., 2015). Unlike these other epithelial appendages, taste papillae are understudied regarding the roles of underlying mesenchyme in their cell differentiation. Studies on molecular regulation of taste papilla cell differentiation have focused on the roles of signaling pathways in the epithelium, including multiple morphogens/growth factors (Beites et al., 2009, Castillo et al., 2014, Castillo-Azofeifa et al., 2017, Gaillard and Barlow, 2011, Gaillard et al., 2017, Gaillard et al., 2015, Iwatsuki et al., 2007, Liu et al., 2007, Liu et al., 2008, Thirumangalathu and Barlow, 2015, Zhou et al., 2006, Petersen et al., 2011, Prochazkova et al., 2017). It has been reported that the molecular programs in the epithelium determines the epithelial cell fate and the position of fungiform taste papillae at E13.5 (Liu et al., 2004, Zhou et al., 2006) when taste papillae protrude from the tongue dorsum (Kaufman, 1992) and remain their stereotypic locations thereafter (Farbman and Mbiene, 1991, Paulson et al., 1985).

In the present study, we found that at the stage (E12) when epithelial cells are rapidly differentiating, the molecular programs in the tongue epithelium are largely governed by the signals from the underlying mesenchyme. Specifically, knockout of a single gene encoding type I BMP receptor *Alk3* in the neural crest-derived tongue mesenchyme leads to many differentially expressed genes (DEGs) in the tongue epithelium, which in turn leads to an absence of taste papillae. This data indicates the requirement of neural crest-derived tongue mesenchyme in taste papilla cell differentiation under the regulation of ALK3-BMP signaling pathway.

Regarding the factors that mediate mesenchymal-epithelial interactions, it has been reported that mesenchymal FGF10, a ligand in the FGF signaling pathway, and Follistatin, a BMP antagonist, regulate the pattern and size of taste papillae in a region-specific manner (Petersen et al., 2011, Prochazkova et al., 2017, Beites et al., 2009). Our data reveal that mesenchyme-specific *Alk3 cKO* does not alter FGF10 and Follistatin expression in the tongue mesenchyme at both transcripts and protein levels. Instead, the production of many proteins is under the control of ALK3-BMP signaling, and proteins within the range of 10-100 kDa regulate taste papilla cell differentiation. The results support that many previously unappreciated proteins from the tongue mesenchyme signal to the overlying epithelium to regulate the cell differentiation.

Regarding the functions of these “novel” mesenchyme-derived proteins, our data indicate two groups of proteins playing opposing roles: promoting or inhibiting taste papilla cell differentiation. Under normal conditions tongue mesenchymal cells secret 10-100 kDa proteins that promote taste papilla cell differentiation. In contrast, proteins (also 10-100 kDa) from *Alk3 cKO* tongue mesenchyme-conditioned medium have profound effects of inhibiting taste papilla formation. Of note, such inhibition of taste papillae occurred in cultured wild type tongues, of which normal mesenchyme was intact and present. Our data reveal that the “inhibitory” proteins from *Alk3 cKO* tongue mesenchyme are potent enough to overwrite the effects of promotants from wild type mesenchyme. The absence of taste papillae in *Alk3 cKO* tongue indicate the necessity of ALK3-BMP signaling to serve as a “brake” inhibiting the production of these “inhibitory” proteins. Further, constitutive activation of ALK3 does not enhance taste papilla cell differentiation support that the normal ALK3-BMP activity is sufficient to inhibit the production of inhibitory proteins. Together, our data support that ALK3-BMP signaling in the tongue mesenchyme, most likely via pSmad1/5/8 (Bragdon et al., 2011, Wang et al., 2014), promotes taste papilla cell differentiation through governing the production of secretory proteins, i.e., enhancing the promotants while suppressing the inhibitors.

Regarding how ALK3-BMP signaling regulates the production of secretory proteins, it is important to note that *Alk3 cKO*-induced alterations of transcripts in the tongue mesenchyme and of proteins in the *Alk3 cKO* mesenchyme-conditioned medium did not overlap. We are aware that the concerns regarding the need for technical optimization cannot be excluded. It is plausible to speculate that ALK3-BMP signaling in the mesenchyme affects the protein production at post-transcriptional levels (e.g., translational regulation, post-translational modifications, secretions).

It is important to note that homogeneous epithelial cells at E11-E11.5 undergo rapid differentiation to the taste papilla placodes and non-gustatory cells between papillae (Barlow, 2015, Kramer et al., 2019). At E11.0-E11.5, all epithelial cells in the lateral tongue swellings are Shh^+^ Krt8^+^. Within a short time period (∼12 hr), the cells either remain Shh^+^ Krt8^+^ clusters and adopt a gustatory/taste papilla cell fate, or become Shh^-^ Krt8^-^ non-gustatory cells (Kramer et al., 2019). Our findings clearly indicate that the mesenchyme via ALK3-BMP signaling promote the gustatory while inhibits the non-gustatory cell fate of the tongue epithelium. Furthermore, in the cultures of E12 wild type tongues of which the epithelial cells have already acquired their gustatory or non-gustatory cell fate, *Alk3 cKO* tongue mesenchyme-derived proteins are potent enough in altering the epithelial cell fate.

### Mesenchymal BMP signaling regulates taste papilla cell differentiation in a receptor-specific manner

The importance of BMP signaling in regulating taste papilla cell differentiation has been documented (Ishan et al., 2020, Beites et al., 2009, Zhou et al., 2006). However, the mediating receptors and involved signals from specific tissue compartments are not clear. In the present study, we detected the four types of type I BMP receptors at different transcripts levels (*Alk3* >> *Alk2* > *Alk6* ≈ *Alk1*) with the *Alk3* being expressed far more rigorously than the other three types. Further phenotypic analyses using mesenchyme-specific knockout of *Alk3* and *Alk2* demonstrated that the BMP signaling mediated by ALK3, not ALK2, is essential for taste papilla cell differentiation, revealing receptor-specific roles of BMP signaling in the tongue mesenchyme in regulating tongue epithelial cell differentiation.

Moreover, p-Smad1/5/8 signals in the tongue mesenchyme and epithelium are robustly detected as early as at E12.0, earlier than the reported starting point (E14.5) in previous studies (Kawasaki et al., 2012). In addition, previous reports (Yumoto et al., 2013, Liu et al., 2018) have shown that taste papillae form after the loss-of-function of non-canonical (pSmad1/5/8-independent) BMP signaling. These data indicate that ALK3-BMP signaling regulates taste papilla cell differentiation most likely via pSmad1/5/8.

The effects of BMPs on taste papilla formation are stage-specific, i.e., promoting taste papilla cell differentiation in rat tongue cultures starting at E13 (≈ mouse E11.5) while inhibiting at E14 (≈ mouse E12.5) (Zhou et al., 2006). The absence of taste papillae in mesenchymal *Alk3 cKO* mice is consistent with (1) the promoting effects of BMP ligands (BMP 2, 4, 7) on taste papilla development at the early embryonic stage and (2) the inhibitory roles of BMP antagonist Follistatin (Beites et al., 2009). It is intriguing that another BMP antagonist Noggin promotes taste papilla cell differentiation in both E13 and E14 rat tongue cultures (Zhou et al., 2006). It is possible that Noggin exert its roles independently of conventional BMP signaling (Bernatik et al., 2017), or that in the 2-day tongue cultures Noggin’s promoting effect that occurred later overwrites its effects at the initial earlier stage. A delicately designed experiment is needed to test which of the possibilities is true. As for the inhibitory effect of BMP ligands on taste papilla formation at the later stage (rat E14), further studies are needed for a clear understanding of the mediating signaling cascades.

The mesenchymal cells in the tongue are largely derived from the neural crest (Liu et al., 2012, Thirumangalathu et al., 2009). It has been reported that *Alk3* deletion may cause the death of neural crest cells immediately after a normal cell migration to reach the destiny (e.g., dorsal aorta) (Morikawa et al., 2009). To understand whether deficient taste papilla cell differentiation in *Alk3 cKO* is caused by the lack of tongue mesenchyme, we analyzed the phenotypic changes at early embryonic stages. At E10.5 when the neural crest-derived cells massively populate the primordia of tongue organ (Han et al., 2012, Ishan et al., 2020), the tongue primordia (i.e., branchial arches I-IV) developed in *Alk3 cKO* similarly to the littermate control. Furthermore, both *in vivo* and *ex vivo* analyses demonstrated that tongue mesenchymal cells are highly proliferating and not apoptotic in *Alk3 cKO*. Our data support that the absence of taste papilla cells in the epithelium is not due to the missing of mesenchyme, but rather is caused by the products of the mesenchyme which are the secretory proteins as discussed above.

### Wnt/β-catenin signaling in tongue epithelium is a downstream target of mesenchymal ALK3-BMP signaling in regulating taste papilla cell differentiation

Wnt/β-catenin pathway is essential for taste papilla development (Liu et al., 2007, Iwatsuki et al., 2007, Zhu et al., 2014, Thirumangalathu and Barlow, 2015). Without Wnt ligands, cytoplasmic β-catenin is constitutively degraded by the destruction complex composed of Axin, adenomatosis polyposis coli (APC), protein phosphatase 2A (PP2A), casein kinase 1α (CK1α) and glycogen synthase kinase 3β (GSK3β) (Komiya and Habas, 2008). Phosphorylation of β-catenin by CK1α and GSK3β leads to ubiquitin-mediated proteolytic destruction, thus making it unavailable for nuclear translocation (Komiya and Habas, 2008). Binding of Wnt ligands to the receptor complex composed of the Frizzled (Fzd) and the lipoprotein receptor-related protein (LRP)5/6 promotes Axin translocation to the cytoplasmic tail of LRP5/6 and activates disheveled (Dvl) and, in turn, inhibits GSK3β, thereby β-catenin proteins may be stabilized and translocated to nucleus to serve as co-transcription factor of Lef1/Tcf (Komiya and Habas, 2008) and/or Prox1 (Liu et al., 2015) in regulating target gene expression.

In mesenchymal *Alk3 cKO*, expression of Wnt/β-catenin signaling components in the epithelium is altered including the down-regulation of 3 encoding genes of key positive regulators (*Fzd10*, *Lgr5, Prox1*) and the up-regulation of the inhibitor *Sostdc1* (Sclerostin domain containing 1) (Lintern et al., 2009). Addition of GSK3β inhibitor LiCl (Iwatsuki et al., 2007) to activate Wnt/β-catenin pathway rescues taste papillae in *Alk3 cKO* tongue cultures indicating that down-regulated Wnt/β-catenin signaling is the main cause of taste papilla loss in *Alk3 cKO*. Noteworthy, we did not detect altered transcripts and protein levels of Wnt/β-catenin agonists and antagonists in *Alk3 cKO* compared to control. Moreover, addition of Wnt3a to the culture medium did not rescue taste papilla formation in *Alk3 cKO* while effectively promoted taste papilla formation in control. Together, our data indicate a defected epithelial Wnt/β-catenin pathway upstream to GSK3β in *Alk3 cKO*, which may be due to the lack of receptors Lgr5 and Fzd10, and the increase of inhibitor Sostdc1.

In summary, our findings bring forward a novel concept to the field of taste biology that during embryonic development, taste papilla cell differentiation requires mesenchymal ALK3-BMP signaling in suppressing the production of inhibitory secretory proteins, thus allow for a proper activity of epithelial Wnt/β-catenin signaling and taste papilla development. It will be important to define the roles of mesenchymal stromal cells and cell products in regulating taste cell renewal of mature taste buds. If this new concept is tested to be true in adults, identification of these regulatory proteins may be beneficial to developing novel therapeutic treatments for taste disorders caused by deficiencies of taste cell differentiation.

## Materials and Methods

### Animal use and tissue collection

The use of animals was approved by the Institutional Animal Care and Use Committee at the University of Georgia. The studies were performed in compliance with the National Institutes of Health Guidelines for the care and use of animals in research. The animals were maintained in the animal facilities in the Department of Animal and Dairy Science at the University of Georgia.

Wild type (WT, C57BL/6J, stock #000664) and <u>n</u>uclear td<u>T</u>omato <u>n</u>uclear E<u>G</u>FP (nTnG) double reporter [(B6; 129S6-*Gt-GT(ROSA)26Sor^tm1(CAG-tdTomato*, EGFP*) Ees^*/J, stock #023035] mice were purchased from the Jackson Laboratory. *Alk2 floxed* (*Alk2^fx/fx^*^)^ (Mishina et al., 1995) and *Alk3 floxed* (*Alk3^fx/fx^*) (Mishina et al., 1995) mice, and mice carrying a constitutively activated form of *Alk3* (*CAG-Z-EGFP-caAlk3*) transgene (hereafter *caAlk3*) (Komatsu et al., 2013) were provided by Dr. Yuji Mishina, University of Michigan. Heterozygous *Wnt1-Cre* [B6.Cg-^Tg (Wnt1-Cre) 11Rth^ Tg(Wnt1-GAL4) 11Rth/J, Jackson Laboratory, Stock# 003829] or *Sox10-Cre* [B6; CBA-Tg (Sox10-Cre) 1Wdr/J, Stock#025807] mice were bred with homozygous *Alk2^fx/fx^* or *Alk3^fx/fx^* mice to generate *Wnt1-Cre/Alk2^fx/+^*, *Wnt1-Cre/Alk3^fx/+^* and *Sox10-Cre/Alk3^fx/+^* mice that were backcrossed with *Alk2^fx/fx^* and *Alk3^fx/fx^* mice to generate conditional knockout (cKO) embryos (*Wnt1-Cre/Alk2 cKO*, *Wnt1-Cre/Alk3 cKO*, and *Sox10-Cre/Alk3 cKO*). To generate *Wnt1- Cre/caAlk3* mice, heterozygous *Wnt1-Cre* mice were bred with homozygous *caAlk3* mice. *Cre^-^* littermates (*Cre^-^/Alk2^fx/fx^*, *Cre^-^/Alk3^fx/fx^* or *Cre^-^/caAlk3*) were used as controls.

Noon of the day on which the dam was positive for a vaginal plug was designated as embryonic day (E) 0.5. Timed pregnant mice were euthanized with CO_2_ followed by cervical dislocation. Embryos (E10.5-E14.5) were dissected from uterus under a dissection microscope. The stages of embryos were confirmed by the number of somite pairs and the development of multiple organs.

The following primers were used for genotyping: 5’- CCCCCATTGAAGGTTTAGAGAGAC - 3’ and 5’- CTAAGAGCCATGACAGAGGTTG -3’ for *Alk2 floxed* (160 bp) and WT (250 bp) fragments; 5’- GCAGCTGCTGCTGCAGCCTCC -3’ and 5’- TGGCTACAATTTGTCTCATGC -3’ for *Alk3 floxed* (230 bp) and WT (150 bp) fragments; 5’- GTGCTGGTTATTGTGCTGTCTC -3’ and 5’- GACGACAGTATCGGCCTCAGGAA -3’ for *caAlk3* gene product (580 bp); 5′- ATTGCTGTCACTTGGTCGTGGC -3′ and 5’- GGAAAATGCTTCTGTCCGTTTGC -3’ for the *Cre* gene product (200 bp).

### Immunohistochemistry on whole-mount organs

Embryonic (E10.5-E14.5) tongues and E12.0+2-day tongue organ cultures were processed for immunohistochemistry as previously described (Liu et al., 2004). Briefly, tongue organs or cultures were fixed in 4% paraformaldehyde (PFA) at 4°C for 2 hr and then washed in 0.1 M phosphate-buffered saline (PBS). Blocking of endogenous hydrogen peroxidase was performed using 6% H_2_O_2_ in methanol followed by an antigen retrieval process, i.e., heating at 95°C for 5 min in antigen retrieval solution (CTS045, R&D Systems, Minneapolis, MN). After blocking non-specific binding in 2% non-fat milk in 0.1 M PBS containing 0.1% Triton X-100 (PBS/MT; X-100, Sigma Aldrich, St. Louis, MO), the organs were incubated in goat anti-Shh primary antibody (Table 1) in 10% normal donkey serum (NDS; D9663, Sigma Aldrich, St Louis, MO) in PBS/MT at 4°C for 48 hr. After thorough washing in PBS/MT on ice (1 hr x 5), the organs were incubated in biotin-conjugated secondary antibody (1:500; BA-5000, Vector Laboratories, Burlingame, CA) in 1% NDS in PBS/MT at 4°C for 24 hr and subjected to peroxidase-conjugated streptavidin treatment in blocking solution (1:500; PK6200, Vector Laboratories, Burlingame, CA) at 4°C for 24 hr after rinsing in PBS/MT on ice (1 hr x 5). Following rinses in PBS/MT (1 hr x 5) and PBT (0.1 M PBS, 0.1% Triton X-100, and 0.2% bovine serum albumin; 1 hr x 2) on ice, tongue organs were processed for DAB (SK4100, Vector Laboratories, Burlingame, CA) pre-incubation (without H_2_O_2_) and reaction (with H_2_O_2_). The immunostained tongues were rinsed in 0.1 M PBS and photographed using an SZX16 Olympus Stereomicroscope.

**Table 1.** Primary antibodies were used for immunohistochemistry.

| Primary antibody | Source (catalog number, company) | Dilution |
| --- | --- | --- |
| Rat anti-BrdU | MCA2060, Bio Rad, Hercules, CA | 1:500 |
| Rabbit anti-c-Cas3 | #9661, Cell Signaling, Danvers, MA | 1:500 |
| Goat anti-E-cadherin | AF748, R&D Systems, Minneapolis, MN | 1:1000 |
| Sheep anti-FGF10 | AF6224, R&D Systems, Minneapolis, MN | 1:10,000 |
| Goat anti-Follistatin | AF669, R&D Systems, Minneapolis, MN | 1:10,000 |
| Rat anti-Krt8 | TROMA-I, Developmental Studies Hybridoma Bank,<br>Iowa city, IA | 1:1000 |
| Sheep anti-Ki67 | AF7649, R&D Systems, Minneapolis, MN | 1:500 |
| Goat anti-Noggin | AF719, R&D Systems, Minneapolis, MN | 1:10,000 |
| Rabbit anti-p-H3 | #13820, Cell Signaling, Danvers, MA | 1:500 |
| Rabbit anti-p-Smad1/5/8 | #13820, Cell Signaling, Danvers, MA | 1:500 |
| Goat anti-Shh | AF464, R&D Systems, Minneapolis, MN | 1:300 |
| Chicken anti-Vimentin | AB5733, EMD Millipore, Burlington, MA | 1: 500 |

### Immunohistochemistry on cells and sections

E11.5 mesenchymal cells or E12.0-E12.5 tongue tissues were fixed in 4% PFA in 0.1 M PBS at 4°C for 15 min (cells) or 2 hr (tongues). PFA-fixed tongue tissues were cryoprotected in 30% sucrose in 0.1 M PBS at 4°C for at least 24 hr, embedded in O.C.T. compound (#23730571, Fisher Scientific, Waltham, MA), and rapidly frozen. Cryostat sections were cut at 10 μm in thickness for immunohistochemistry.

Cells or tongue sections were air-dried at room temperature for 1 hr, then rehydrated in 0.1 M PBS (5-10 min x 3). Blocking of nonspecific binding was carried out by incubation with 10% NDS in 0.1 M PBS containing 0.3% Triton X-100 at room temperature for 30 min. Then the cells or sections were incubated with primary antibodies (Table 1) in carrier solution (1% normal donkey serum, 0.3% Triton X-100 in 0.1 M PBS) at 4°C for overnight. Cells and sections treated without a primary antibody were used as negative controls. After rinsing in 0.1 M PBS (5-10 min x 3) at room temperature, cells or sections were incubated with Alexa Fluor® 488 or 647-conjugated secondary antibody (1:500; Invitrogen, Eugene, OR) in carrier solution at room temperature for 1 hr, rinsed with 0.1 M PBS (5-10 min x 3) at room temperature and then counterstained with DAPI (200 ng/ml in PBS; D1306, Life Technologies, Carlsbad, CA) at room temperature for 10 min. After thorough rinsing in 0.1 M PBS, cells or sections were air-dried and mounted with Prolong^®^ Diamond antifade mounting medium (P36970, Fisher Scientific, Waltham, MA). Immunostained sections and cells were examined under a fluorescent light microscope (EVOS FL, Life Technologies, Carlsbad, CA). A laser scanning confocal microscope (Zeiss LSM 710, Biomedical Microscopy Core at the University of Georgia) was used to take single-plane images.

### Immunohistochemistry on epithelial sheets

The tongue epithelium was separated from the mesenchyme in postnatal day 21 (P21) *Wnt1-Cre/caAlk3* and *Cre^-^* littermate control mice as previously described (Liu et al., 2008, Venkatesan et al., 2016). Separated epithelial sheets were immunostained for pan taste cell marker Krt8 (Table 1) as previously described (Liu et al., 2008, Ishan et al., 2020, Venkatesan et al., 2016) and photographed using an SZX16 Olympus Stereomicroscope.

### *In situ* hybridization

Tongues were dissected from E12.0 *Wnt1-Cre/Alk3 cKO* and *Cre^-^/Alk3^fx/fx^* littermate control embryos in 0.1 M PBS and fixed with 4% PFA at 4°C for 24 hr. PFA-fixed tissues were processed for cryostat sectioning at 15 μm in thickness. Conventional *in situ* hybridization for *Alk3* was performed as previously described (Lauter et al., 2011) using digoxigenin-labeled riboprobes. Plasmids carrying the *Alk3* probe was provided by Dr. Yuji Mishina (University of Michigan). Digoxigenin-labeled antisense *Alk3* RNA probe was prepared by linearization with EcoR1 (New England Biolabs, Ipswich, MA) and transcription with T7 RNA polymerase (Promega, Madison, WI).

For RNAscope® *in situ* hybridization, RNAscope® Intro Pack 2.5 HD Reagent Kit Brown (#322300, Advanced Cell Diagnostics, Newark, CA) was used on E12.0 WT tongue sections. *Alk2* (# 312411) and *Alk3* (# 312421) probes were purchased from the Advanced Cell Diagnostics and *in situ* hybridization was performed following the manufacturer’s instructions. Cell nuclear counterstaining was performed with 50% Hematoxylin.

### Scanning electron microscopy (SEM)

E10.5-E11.5 *Cre^-^/Alk3^fx/fx^* littermate control and *Wnt1-Cre/Alk3 cKO* mutant branchial arches (BAs) or tongues were fixed in 2.5% glutaraldehyde (#75520; Electron Microscopy Science, Hatfield, PA) and 4% PFA in 0.1 M PBS (pH 7.3) at 4°C for 24 hr. After rinsing in 0.1 M PBS at room temperature (10 min x 3) and post-fixed in a sequence of aqueous 1% O_S_O_4_ (#19150, Electron Microscopy Science, Hatfield, PA) in 0.1 M PBS, 1% tannic acid (#16201, Sigma Aldrich, St. Louis, MO) in MQ-H_2_O, and 1% O_S_O_4_ in MQ-H_2_O, on ice for 1 hr each, tissues were dehydrated in an ascending series of ethanol (35, 50, 70, 90, 100%), and hexamethyldisilazane (HMDS, #440191; Sigma Aldrich, St. Louis, MO) at room temperature (1 hr x 3). After a slow air dry in a fume hood, tissues were mounted on specimen stubs and sputter-coated with gold/palladium (Leica Gold/Carbon coater; Georgia Electron Microscope Core Facility, University of Georgia). Tissues were then imaged using a scanning electron microscope (FEI Teneo FE-SEM; Georgia Electron Microscope Core Facility, University of Georgia).

### Collection of conditioned medium from mesenchymal cell cultures

The tongue mesenchyme was separated from the epithelium in E11.5 *Wnt1-Cre/Alk3 cKO* and *Cre^-^/Alk3^fx/fx^* littermate control mice as previously described ^(Liu et al., 2008)^. Separated mesenchyme was then transferred to a culture dish, cut into small pieces, and cultured in a humidified CO_2_ incubator at 37°C in a serum-free medium, i.e., a 1:1 mixture of Dulbecco’s modified Eagle’s medium and Ham’s nutrient F12 (#11320033, DMEM/F12, GIBCO, Gaithersburg, MD) containing 50 μg/ml gentamicin sulfate (#15750060, GIBCO, Gaithersburg, MD). After 1 day in cultures, the medium was replaced with fresh medium and continued to culture for 2 days. Culture medium was collected as conditioned medium.

### Extraction of proteins and protein fractions from conditioned medium

Proteins were extracted from the conditioned medium using the protein precipitation kit (#2100, Millipore Sigma, Burlington, MA) following the manufacturer’s specifications. To isolate proteins at different molecular weights, mesenchyme-conditioned media were filtered through 100-kDa followed by 10-kDa Amicon® filters (UFC910024, UCF910008, Millipore Sigma, Burlington, MA) by centrifugation at 4000g for 10 min to obtain >100-kDa and 10 to 100-kDa proteins as described (Whittaker et al., 2020). Proteins (<10-kDa) from the medium leaked through 10-kDa filters were extracted using the protein precipitation kit as aforementioned. The supernatant was used as the residual solution.

### Liquid chromatography-mass spectrometry analysis of proteins in mesenchyme-conditioned medium

Proteins isolated from the *Wnt1-Cre/Alk3 cKO* and *Cre^-^/Alk3^fx/fx^* conditioned medium were reduced with 5 mM of tris (2-carboxyethyl) phosphine hydrochloride, alkylated with 13.75 mM of iodoacetamide, and digested with trypsin/Lys-C mix (V5071, Promega, Madison, WI). The resulting peptides were cleaned up with Acclaim™ PepMap™ 100 C18 spin columns (SEM SS18V, The Nest Group, Ipswich, MA) dried down, and reconstituted in 0.1% formic acid. The reconstituted peptides were separated on an Acclaim™ PepMap™ 100 C18 column and eluted into the nano-electrospray ion source of an Orbitrap Eclipse™ Tribrid™ mass spectrometer (Thermo Fisher Scientific, Waltham, MA) at a flow rate of 200 nl/min. The elution gradient consists of 1-40% acetonitrile in 0.1% formic acid over 220 min followed by 10 min of 80% acetonitrile in 0.1% formic acid. The spray voltage was set to 2.2 kV and the temperature of the heated capillary was set to 275°C. The full mass spectrometry scans were acquired from m/z 300 to 2000 at 60k resolution in the orbitrap, and the MS2 scans for the most intense precursors were fragmented via collision-induced dissociation (CID) and collected in the ion trap.

The raw spectra were searched against mouse protein database (UniProt) by SEQUEST using Proteome Discoverer (v2.5, Fisher Scientific, Waltham, MA). The mass tolerance was set as 20 ppm for precursors and 0.5 Da for fragments. The peptide-spectrum matches (PSMs) resulted from the database search were identified, quantified, and filtered to a 10% peptide false discovery rate (FDR) and then clustered into a final protein-level FDR of 1%. The NSAF (Normalized spectral abundance factor) values were calculated for each protein and used to quantify their relative abundance and fold change across samples.

Gene Ontology (GO) enrichment and pathway analysis (http://www.geneontology.org/GO.database.shtml) was done to analyze the top 10 functional associations of the identified differentially expressed proteins (control only or *Alk3 cKO* only with ≥ 100 NSAF) from the mass spectrometry analyses.

### Tongue organ cultures

E12.0+2-day tongue organ cultures were performed as previously described (Mbiene et al., 1997, Mistretta et al., 2003). To test the impacts of tongue mesenchyme on the epithelial cell differentiation and taste papilla formation, the mesenchymal tissue or mesenchyme-conditioned medium of E11.5-12.0 *Cre^-^/Alk3^fx/fx^* or *Wnt1-Cre/Alk3 cKO* tongues was added to E12.0 tongue cultures from wild type mice. Proteins from conditioned medium (>100 kDa, 10-100 kDa, or <10 kDa at a concentration of 200 µg/ml), or residual solution was added to the standard culture medium.

To activate Wnt/β-catenin signaling, 5 mM LiCl (Clément-Lacroix et al., 2005) or 20% Wnt3a conditioned medium (J2-001, MBL international, Woburn, MA) were added to the culture medium. To digest proteins from conditioned medium, an equal amount of Proteinase K (#3115879001, Sigma Aldrich, St Louis, MO) (i.e., 200 µg/ml Proteinase K for 200 µg/ml of proteins) was added and incubated at 37°C for 6 hr. Proteinase K was inactivated by heating at 95°C for 10 min before the ProK-pretreated proteins were added to culture medium. After 2 days, cultures were collected and processed for analyzing taste papilla formation and epithelial cell differentiation using Shh immunosignals in the tongue cultures.

### RNA sequencing and quantitative reverse transcriptase-polymerase chain reaction (qRT-PCR)

Separated mesenchymal and epithelial tissues from E12.0 *Cre^-^/Alk3^fx/fx^* and *Wnt1-Cre/Alk3 cKO* tongues were immersed in Trizol solution (#15596018, Life Technologies, Carlsbad, CA) for RNA extraction using the RNeasy Plus kit (#74136, Qiagen, Hilden, Germany). For each experimental condition, a total of nine mesenchymal and epithelial tissues (pooled three tissues x three replicates) were used. RNA concentrations were measured using Nanodrop 8000 spectrophotometer (Nanodrop, Thermo Scientific, Waltham, MA).

RNA sequencing was performed in Georgia Genomics and Bioinformatics Core Facility at the University of Georgia using the NextSeq 500 system (Illumina, San Diego, CA) following the procedures as described previously (Ishan et al., 2021). Raw data were mapped to mouse reference genome GRCm38 (mm10) using STAR (Dobin et al., 2013). Transcripts were analyzed and reported as FPKM (fragments per kilobase per million) by StringTie (Pertea et al., 2015). Differentially expressed genes (DEGs) were detected using the R package DESeq2 (Love et al., 2014). GO and Kyoto Encyclopedia of Genes and Genomes (KEGG) pathway analyses were used to investigate the functional associations of the DEGs. R package ggplot2 was used to illustrate data as histograms.

For qRT-PCR analyses, complementary DNA (cDNA) was synthesized from the extracted RNA using SuperScript™ First-Strand Synthesis System (#11902018, Fisher Scientific, Waltham, MA). The expression of the *Alk3* gene was detected using 5’-GCAGCTGCTGCTGCAGCCTCC -3’ and 5’- TGGCTACAATTTGTCTCATGC -3’ primers. Changes in gene expression levels in *Wnt1-Cre/Alk3 cKO* and *Cre^-^/Alk3^fx/fx^* tongue epithelium and mesenchyme are presented as means ± standard deviation (X±SD; n=3) of 2^−ΔCT^ values.

### Western blot

To detect the levels of proteins extracted from the E12.0 tongue mesenchyme and E11.5 mesenchyme-conditioned medium of *Wnt1-Cre/Alk3 cKO* and *Cre^-^/Alk3^fx/fx^* mice, SDS-PAGE and Western blot were conducted as described previously (Ishan et al., 2021). Proteins from the tongue mesenchyme were extracted using Radioimmunoprecipitation assay (RIPA) buffer (1% NP-40, 150 mmol/L NaCl, 50 mmol/L Tris-HCI, 0.5% sodium deoxycholate, 0.1% SDS, 1 mmol/L EDTA, pH 7.4).

### Quantification and statistical analyses

To quantify the number of Ki67^+^, BrdU^+^, p-H3^+^, c-Cas3^+^, and p-Smad1/5/8^+^ cells per unit area (mm^2^) in tongue sections of E12. *Wnt1-Cre/Alk3 cKO* and *Cre^-^/Alk3^fx/fx^* littermate control mice (n=3 each group), serial sections were immunostained for Ki67, BrdU, p-H3, c-Cas3, and p-Smad1/5/8 and thoroughly analyzed under a fluorescent light microscope (EVOS FL, Life Technologies, Carlsbad, CA). Single-plane laser scanning confocal photomicrographs were taken from every other section using a laser scanning confocal microscope (Zeiss LSM 710, Biomedical Microscopy Core at the University of Georgia). Labeled cells were quantified on the sections of the anterior tongue region. For quantification of the number of fungiform papillae in the E12+2-day WT tongue cultures, bright-field images of anti-Shh immunostained tongues were used. To quantify the Vimentin^+^ and Ki67^+^ cells, single-plane laser scanning confocal photomicrographs were taken from the Vimentin and Ki67 immunostained E11.5+3-day cultures of mesenchymal cells from *Wnt1-Cre/Alk3 cKO* and *Cre^-^/Alk3^fx/fx^* tongues. Numbers of Vimentin^+^ and Ki67^+^ cells were counted in relation to the total number of DAPI^+^ cells and presented as a percentage of Vimentin^+^ and Ki67^+^ cells.

Data are presented as means ± standard deviation (X±SD; n=3). Two-way analyses of variance (ANOVA) followed by Fisher’s LSD analyses were used to compare: (a) numbers of fungiform papillae in E12+2-day WT tongue cultures; (b) Ki67^+^, BrdU^+^, p-H3^+^, c-Cas3^+^, and p- Smad1/5/8^+^ cells per unit area (mm^2^) in tongue epithelium and mesenchyme from *Wnt1- Cre/Alk3 cKO* and *Cre^-^/Alk3^fx/fx^* littermate mice; (c) percentage of Vimentin^+^ and Ki67^+^ cells detected from cell cultures of *Wnt1-Cre/Alk3 cKO* and *Cre^-^/Alk3^fx/fx^* tongue mesenchyme. The Student’s *t*-test was used to evaluate the statistical significance of the differences between Western blot band intensities. A *p*-value less than 0.05 was taken as statistically significant.

## Acknowledgments

This study was supported by the National Institute on Deafness and Other Communication Disorders (NIDCD), NIH (R01DC012308) and The University of Georgia Internal Funding to HXL; and by National Institute of Dental and Craniofacial Research, NIH (R01DE020843) to YM

---

**Supplemental Figure 1.**
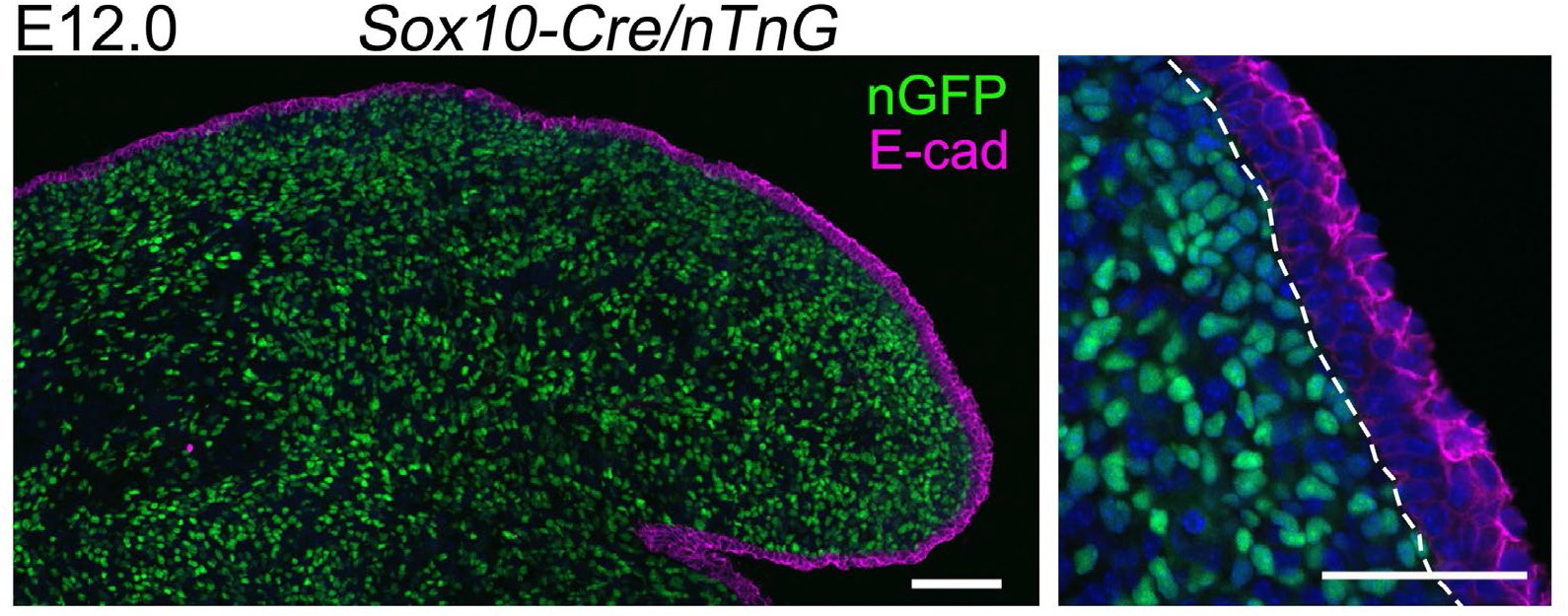
S*o*x10*-Cre*-labeled cells are restricted to the tongue mesenchyme at E12.0.1. Single-plane laser scanning confocal images of sagittal tongue sections from E12.0 *Sox10-Cre/nTnG* mice. The tongue sections were immunostained for epithelial cell marker E-cadherin (E-cad, magenta). The image to the right illustrates the anterior tongue region at a high magnification. Dashed lines demarcate the tongue epithelium from the underlying mesenchyme. Scale bars: 50 μm.

**Supplemental Figure 2.**
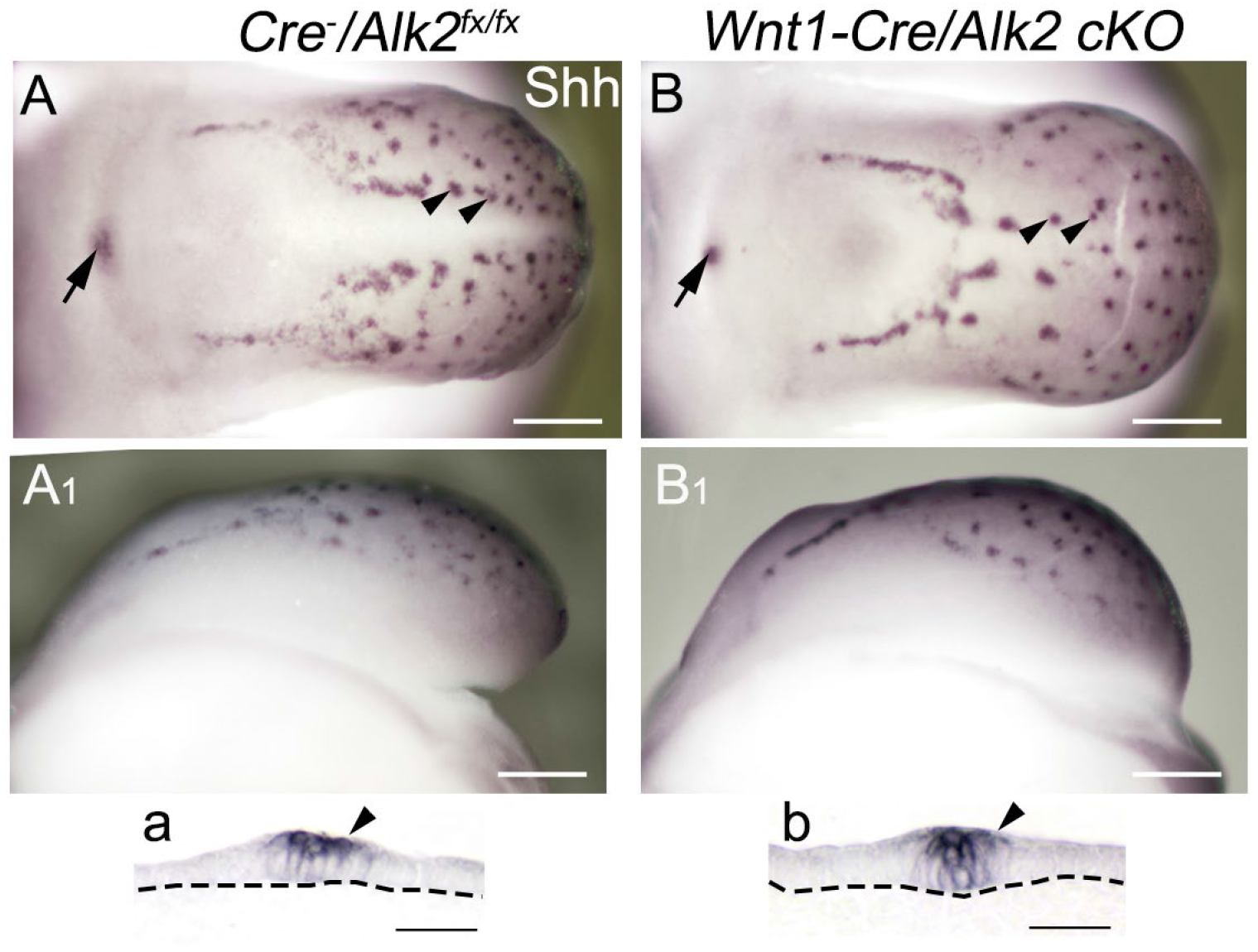
Taste papillae develop in the E12.0 *Wnt1-Cre/Alk2 cKO* mice. Light microscopy images of E12.5 *Cre^-^/Alk2^fx/fx^* (A, A_1_) and *Wnt1-Cre/Alk2 cKO* (B, B_1_) tongues that were immunostained for Shh. A-B: dorsal view; A_1_-B_1_: side view; a-b: sagittal tongue sections of *Cre^-^/Alk2^fx/fx^* (a) and *Wnt1-Cre/Alk2 cKO* (b) mice. Dashed lines demarcate the tongue epithelium from the underlying mesenchyme. Arrowheads and arrows point to Shh^+^ fungiform and circumvallate papilla placodes respectively. Scale bars: 200 µm in A-B, 25 µm in a-b.

**Supplemental Figure 3.**
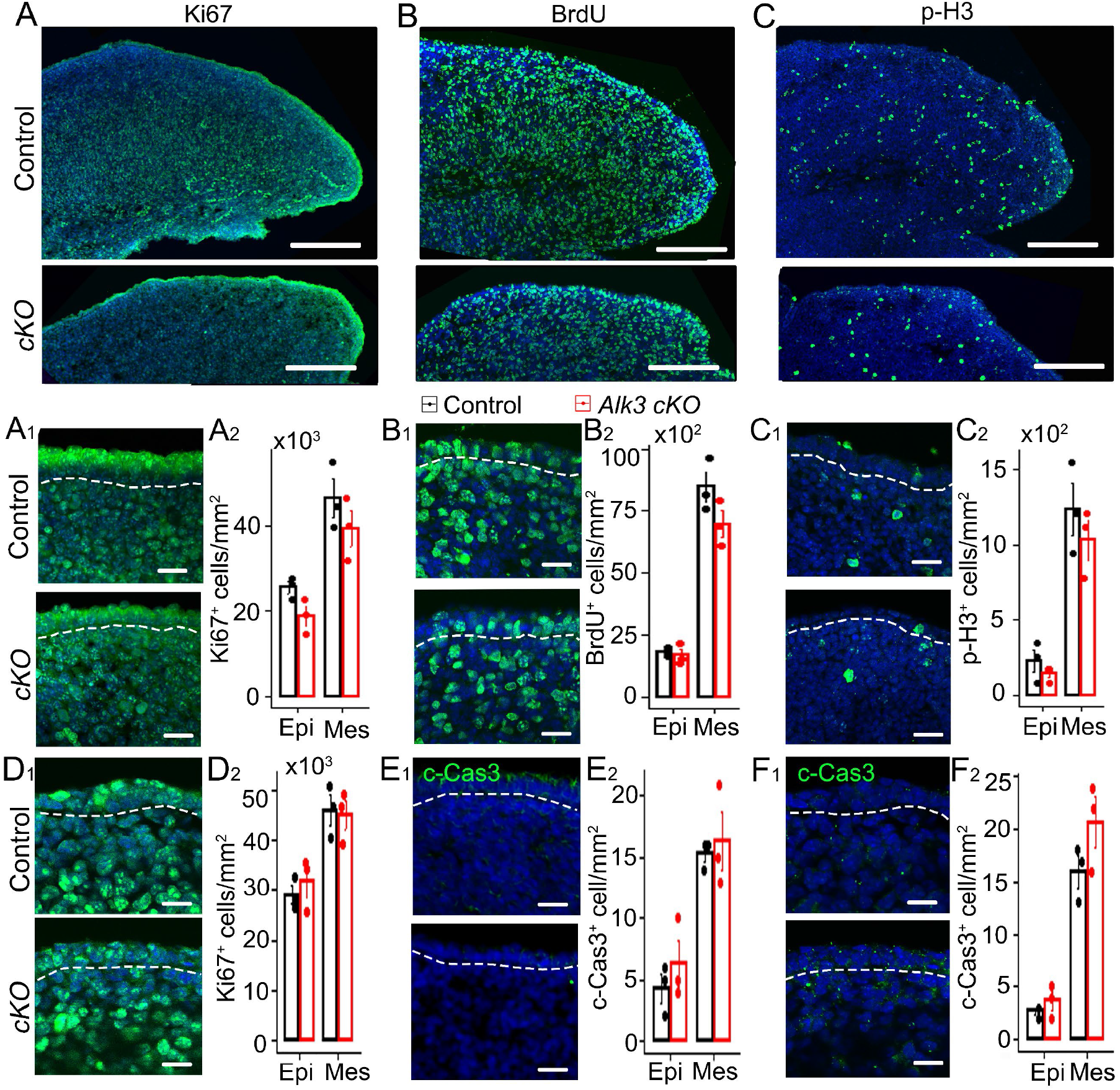
Cell proliferation was unaltered and apoptotic cells were rarely seen in the E12.0 *Wnt1-Cre/Alk3 cKO* or *Sox10-Cre/Alk3 cKO* mouse tongues. **<u>A-F</u>:** Single-plane laser scanning confocal images from E12.0 *Cre^-^/Alk3^fx/fx^* (Control) and *Wnt1-Cre/Alk3 cKO* (*Alk3 cKO* in A-C, E) or *Sox10-Cre/Alk3 cKO* (*Alk3 cKO* in D, F) tongue sections immunostained for cell proliferation marker Ki67 (pan, green in A, D), BrdU (S-phase, green in B), or p-H3 (M-phase, green in C) or apoptosis marker cleaved caspase3 (c-Cas3, green in E, F). **<u>A_1_-F_1_</u>:** High magnification images of the anterior tongue tip. Dashed lines demarcate the tongue epithelium from the underlying mesenchyme. Scale bars: 50 μm. **<u>A_2_-F_2_:</u>** Histograms (X±SD; n=3) to present the number of Ki67^+^, BrdU^+^, p-H3^+^ and c-Cas3^+^ cells per mm^2^ in the epithelium and mesenchyme of *Cre^-^/Alk3^x/fx^* and *Alk3 cKO* tongues. No statistically significant differences were found in *Alk3 cKO* compared to the corresponding regions of *Cre^-^/Alk3^fx/fx^* littermate control using two-way ANOVA followed by Fisher’s LSD analyses.

**Supplemental Figure 4.**
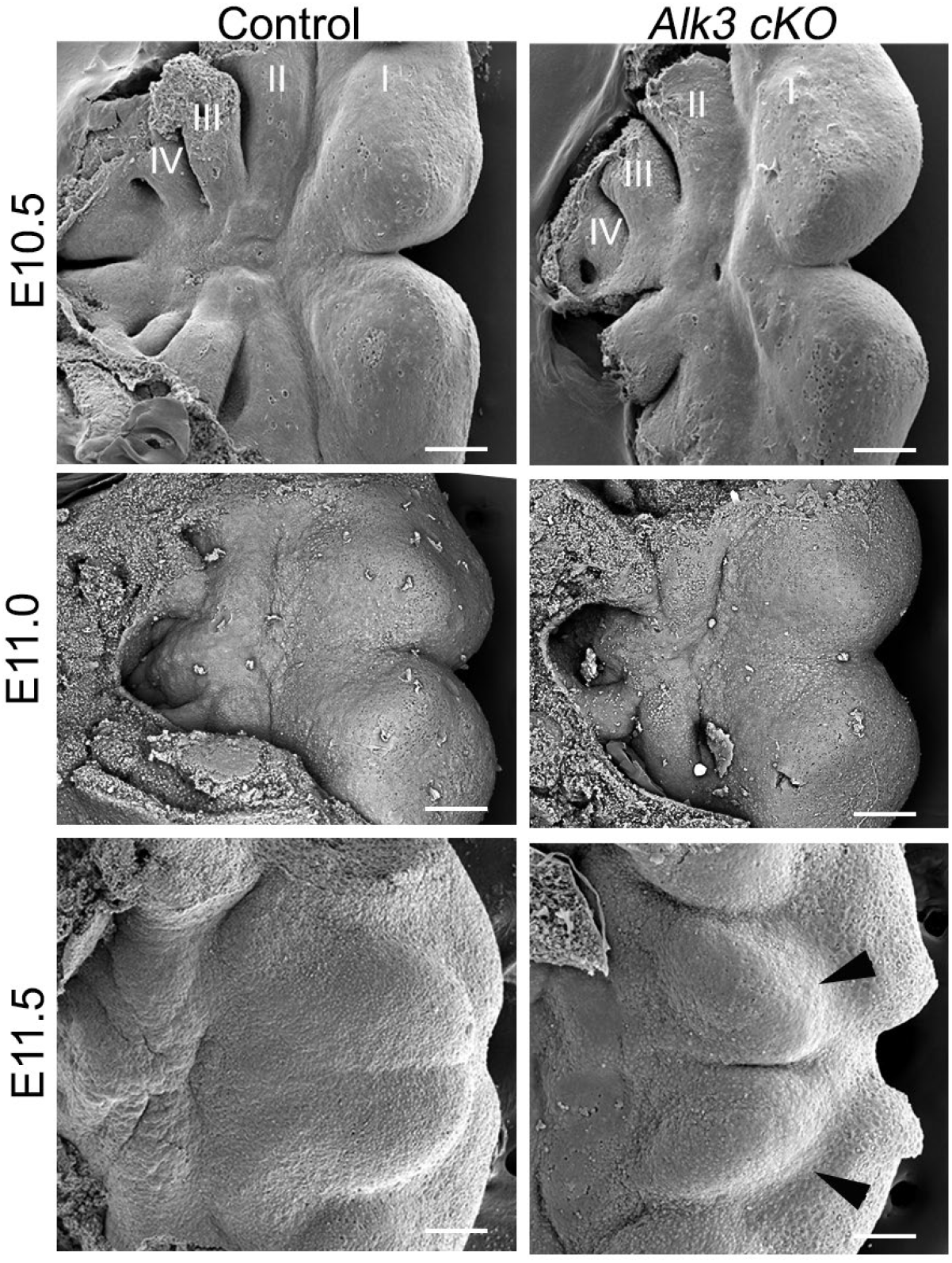
W*n*t1*-Cre/Alk3 cKO* mice do not depict an obvious phenotype of tongue development at early stages until E11.5. Scanning electron microscopy images of E10.5-E11.5 *Wnt1-Cre/Alk3 cKO* (*Alk3 cKO*) and *Cre^-^/Alk3^fx/fx^* littermate (Control) mice. Arrowheads point to lateral tongue swellings. Roman numeral I-IV represents branchial arches 1-4. Scale bars: 100 µm.

**Supplemental Figure 5.**
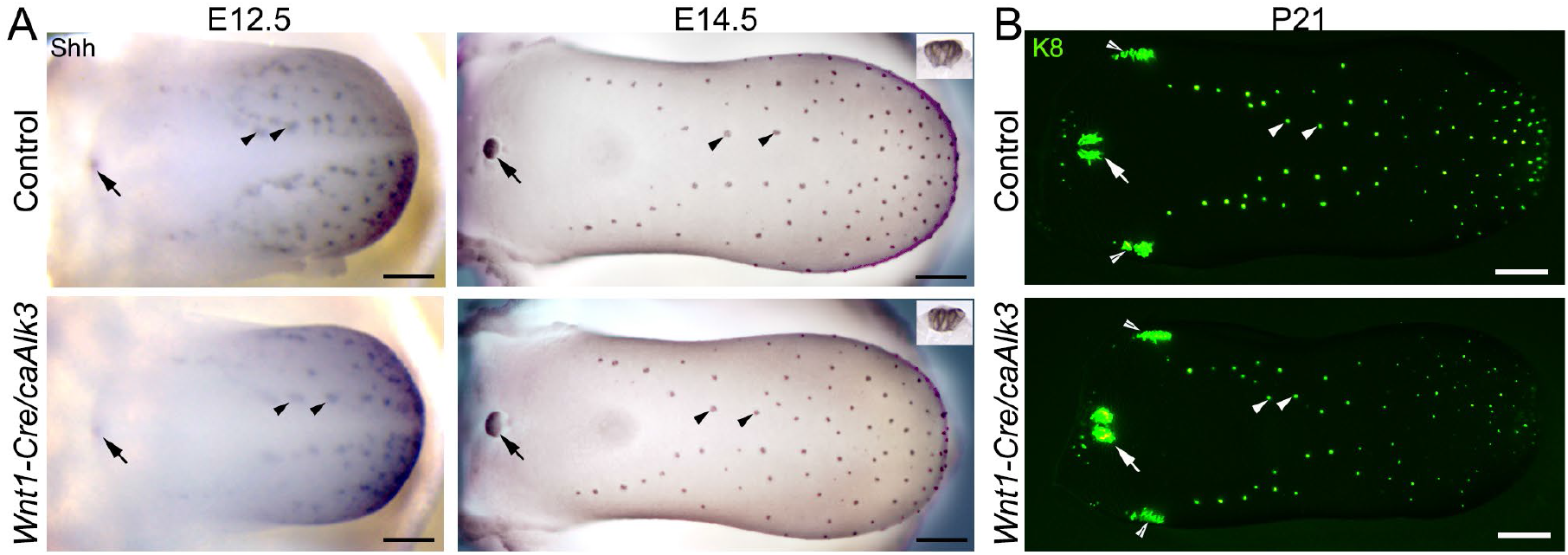
Constitutive activation (*ca*) of *Alk3* in the tongue mesenchyme (*Wnt1-Cre*) does not alter the development of taste papillae and taste buds. **<u>A</u>:** Light microscopy images of E12.5 and E14.5 *Wnt1-Cre/caAlk3* and *Cre^-^* littermate (Control) tongues. Tongues were immunostained for taste papilla marker Shh (blue). Insets are light microscopy images of sagittal tongue sections. Scale bars: 200 µm. **<u>B</u>:** Stereomicroscopy images of postnatal day 21 (P21) control and *Wnt1-Cre/caAlk3* tongue epithelial sheets. Epithelial sheets were immunostained for pan-taste cell marker Krt8. Arrowheads, open arrowheads, and arrows point to fungiform, foliate, and circumvallate taste papillae/buds respectively. Scale bars: 500 µm.

**Supplemental Figure 6.**
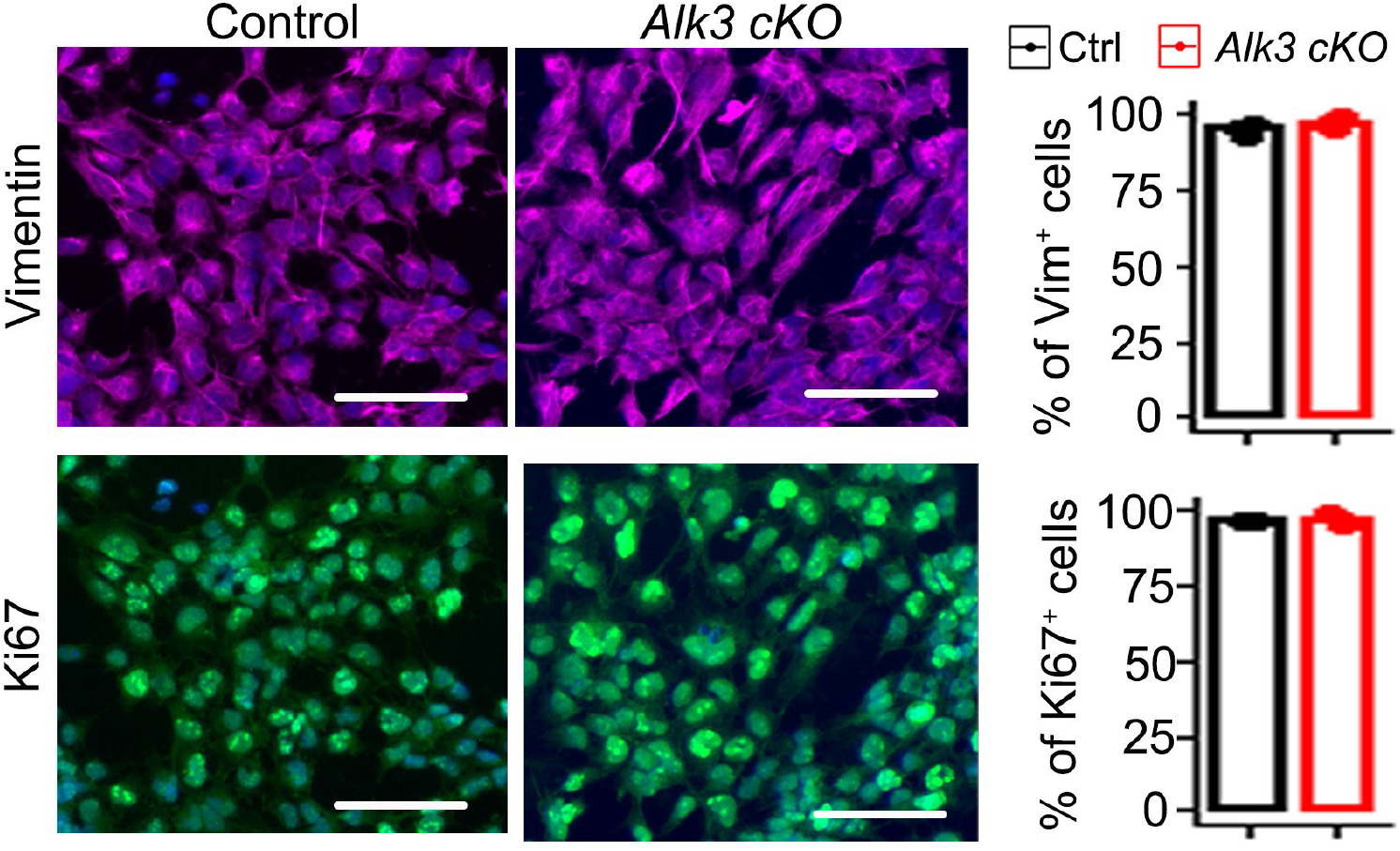
The morphology and proliferating rate were unaltered in the *Wnt1-Cre/Alk3 cKO* tongue mesenchymal cells. Single-plane laser scanning confocal images of E11.5+3-day mesenchymal cell cultures from *Cre^-^/Alk3^fx/fx^* (Control) and *Wnt1-Cre/Alk3 cKO* (*Alk3 cKO*) tongues. Cells were immunostained for mesenchymal cell marker Vimentin (purple) and Ki67 (green), and counter-stained with nuclear marker DAPI (blue). Scale bars: 50 µm. Histograms (X±SD; n=3) on the right present the percentage of Vimentin^+^ or Ki67^+^ cells relative to the total number of cells (DAPI^+^) in the cultures. No statistically significant differences were found in *Alk3 cKO* group compared to the *Cre^-^/Alk3^fx/fx^* littermate control group using two-way ANOVA followed by Fisher’s LSD analyses.

**Supplemental Figure 7.**
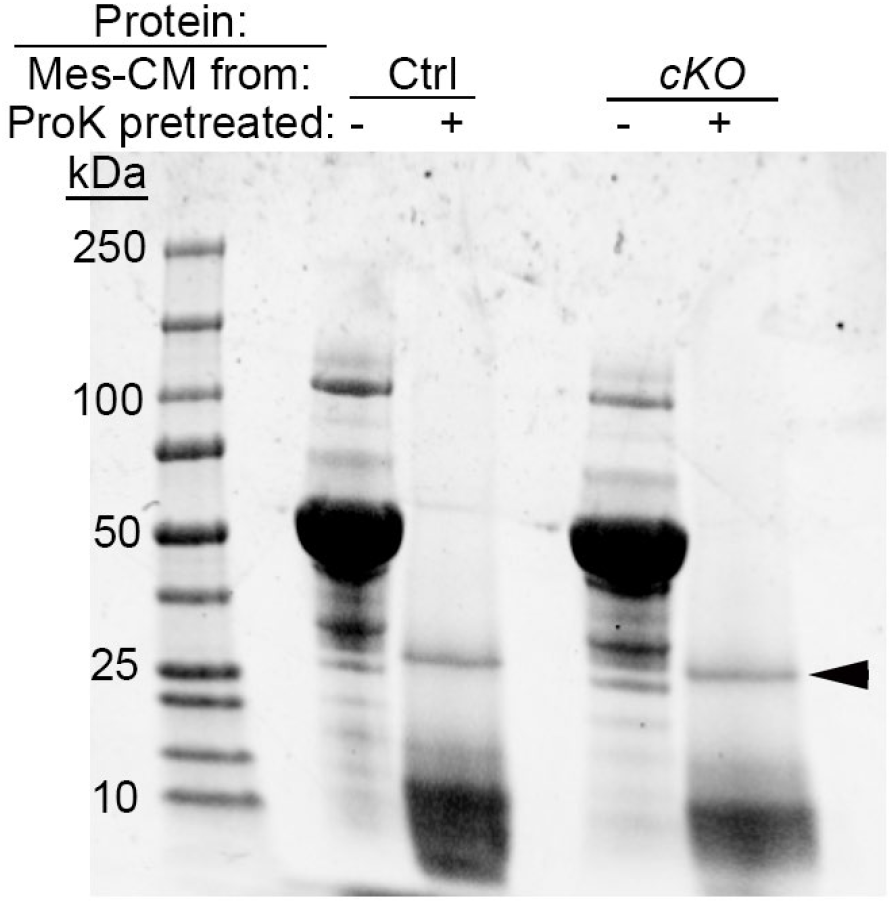
Proteinase K pretreatment efficiently digests the proteins from mesenchyme-conditioned medium. Sodium dodecyl sulfate-polyacrylamide gel image to present the bands of proteins from mesenchyme-conditioned medium (Mes-CM) in *Cre^-^/Alk3^fx/fx^* (Ctrl) and *Wnt1-Cre/Alk3 cKO* (*cKO*) mice without (-) or with (+) proteinase K (ProK) pretreatment. Protein bands are absent after ProK treatment. The arrowhead points to the band of ProK enzyme (28 kDa).

**Supplemental Figure 8.**
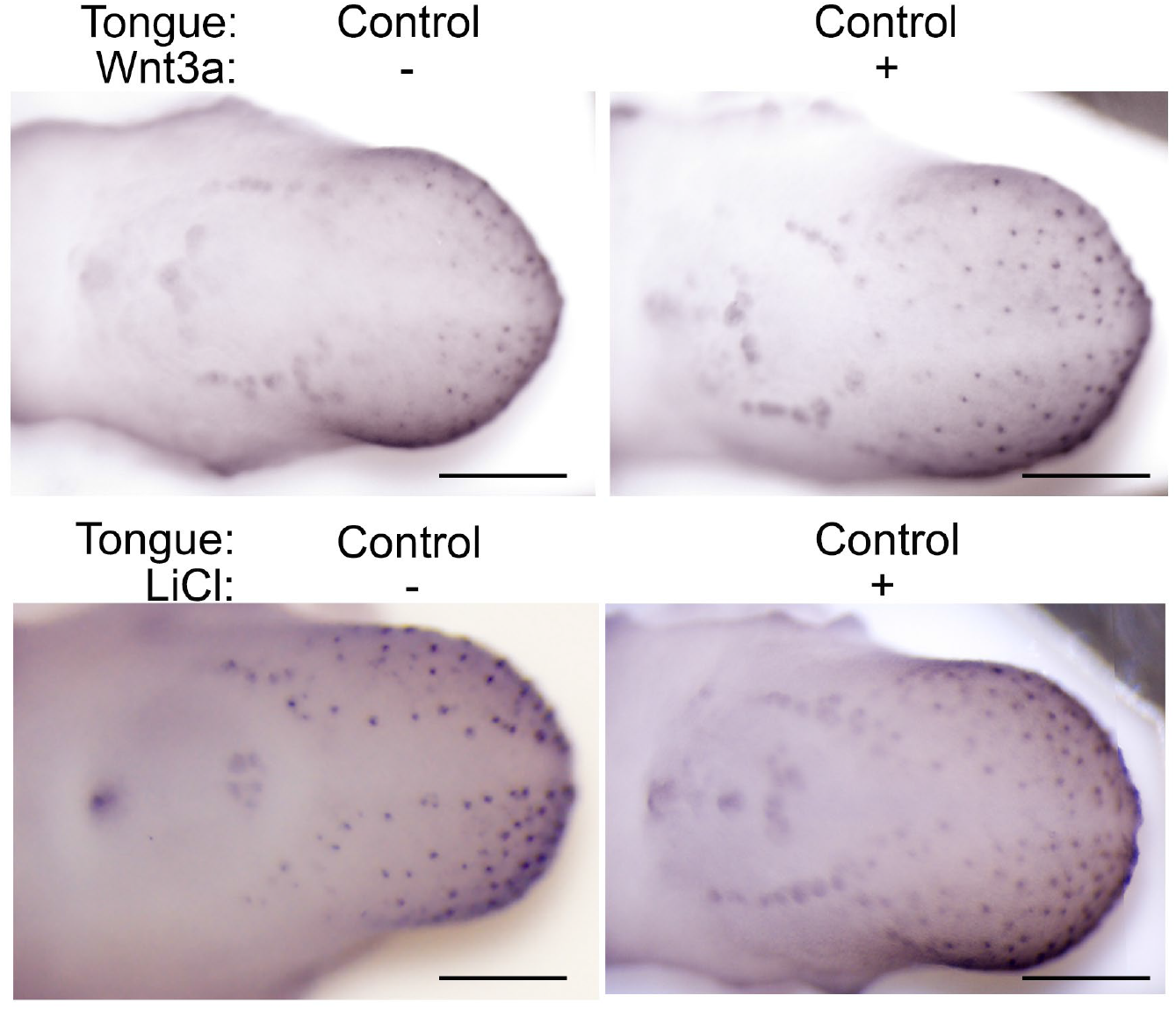
Activation of Wnt/β-catenin signaling promotes taste papilla development in control tongue cultures. Representative light microscopy images of E12+2-day tongue cultures from *Cre^-^/Alk3^fx/fx^* (Control) mice. Tongue cultures were administered with 5 mM LiCl or 20% Wnt3a conditioned medium to activate Wnt/β-catenin signaling activity and immunostained for Shh (blue). Scale bars: 200 µm.

